# Pharmacological inhibition of bromodomain and extra-terminal proteins induces NRF-2-mediated inhibition of SARS-CoV-2 replication and is subject to viral antagonism

**DOI:** 10.1101/2022.09.22.508962

**Authors:** Baxolele Mhlekude, Dylan Postmus, January Weiner, Saskia Stenzel, Francisco J. Zapatero-Belinchón, Ruth Olmer, Jenny Jansen, Anja Richter, Julian Heinze, Nicolas Heinemann, Barbara Mühlemann, Simon Schroeder, Terry C. Jones, Marcel Alexander Müller, Christian Drosten, Andreas Pich, Volker Thiel, Ulrich Martin, Daniela Niemeyer, Gisa Gerold, Dieter Beule, Christine Goffinet

## Abstract

Inhibitors of bromodomain and extra-terminal proteins (iBETs), including JQ-1, have been suggested as potential therapeutics against SARS-CoV-2 infection. However, molecular mechanisms underlying JQ-1-induced antiviral activity and its susceptibility to viral antagonism remain incompletely understood. iBET treatment transiently inhibited infection by SARS-CoV-2 variants and SARS-CoV, but not MERS-CoV. Our functional assays confirmed JQ-1-mediated downregulation of ACE2 expression and multi-omics analysis uncovered induction of an antiviral NRF-2-mediated cytoprotective response as an additional antiviral component of JQ-1 treatment. Serial passaging of SARS-CoV-2 in the presence of JQ-1 resulted in predominance of ORF6-deficient variants. JQ-1 antiviral activity was transient in human bronchial airway epithelial cells (hBAECs) treated prior to infection and absent when administered therapeutically. We propose that JQ-1 exerts pleiotropic effects that collectively induce a transient antiviral state that is ultimately nullified by an established SARS-CoV-2 infection, raising questions on their clinical suitability in the context of COVID-19.

## INTRODUCTION

There is an unmet need for novel effective therapies against the evolving variants of severe acute respiratory syndrome coronavirus-2 (SARS-CoV-2), the causative agent of the coronavirus disease-2019 (COVID-19) pandemic. The hallmark of severe SARS-CoV-2 infections is an excessive inflammatory response resulting in tissue damage and multiorgan failure (Bülow Anderberg et al., 2021). Therefore, therapeutic avenues that simultaneously dampen SARS-CoV-2 replication and antagonise its pathophysiological effects are utmostly desired.

A group of small molecular inhibitors of the bromodomain and extra-terminal (BET) proteins (iBETs) has been suggested to hold potential for the realisation of such therapies (Gilham et al., 2021; Mills et al., 2021; Samelson et al., 2022; Vann et al., 2022). The BET protein family is made up of four multifaceted, ubiquitously expressed and evolutionarily conserved proteins: bromodomain-containing protein (BRD) BRD2, BRD3, BRD4, and BRTD (Lara-Ureña and García-Domínguez, 2021). They act as epigenetic readers and transcriptional co-activators by binding to the acetylated lysine residues in histones in the chromatin and recruit the cellular transcriptional machinery to drive transcription of their target genes (Cheung et al., 2021; Suarez-Alvarez et al., 2017). Alternatively, BET proteins can act as transcriptional co-repressors by interacting with their cellular partners to form repressor complexes, which suppress inappropriate transcriptional programs to maintain tissue homeostasis (Cheung et al., 2021).

The ability of BET proteins to regulate transcription has made them prime targets for several diseases that hijack the cellular transcriptional machinery, including SARS-CoV-2 infection (Chen et al., 2022; De Rijck et al., 2013; Gordon et al., 2020; Shorstova et al., 2021; Wu et al., 2016). iBETs evict BET proteins from histones and their non-histone binding partners (Stathis and Bertoni, 2018). Therefore, clinical development of iBETs, most of which are in different phases of clinical trials in the context of cancer therapy and other pathologies (Hajmirza et al., 2018; Shorstova et al., 2021), may provide a pharmacological tool for therapeutic intervention against several diseases driven by the BET protein-mediated transcriptional programs.

BRD2 co-activates expression of ACE2 (Samelson et al., 2022), the main receptor for SARS-CoV-2 entry into target cells (Hoffmann et al., 2020). Consequently, treatment of target cells with iBETs downregulates ACE2 expression and inhibits SARS-CoV-2 replication and inflammatory responses (Gilham et al., 2021; Mills et al., 2021; Samelson et al., 2022). Additionally, BET proteins co-activate the interferon (IFN) (Patel et al., 2013) and NF-κB (Huang et al., 2009) signalling pathways, which are the main drivers of the inflammatory responses in COVID-19 patients (Lee et al., 2020; Ramasamy and Subbian, 2021).

Despite the growing literature on the anti-SARS-CoV-2 activity of iBET candidates (Gilham et al., 2021; Mills et al., 2021; Samelson et al., 2022; Vann et al., 2022), key questions remain regarding the genome-wide epigenetic alterations orchestrating iBET-mediated transcriptional responses that underlie their anti-SARS-CoV-2 activity. The missing information about the susceptibility of iBET-mediated antiviral activity to SARS-CoV-2 subversion is hampering their potential as the next generation of prophylactics in the context of COVID-19. Here, we sought to address these key questions by conducting an in-depth functional and multi-omics analysis of the signalling alterations that underlie JQ-1-mediated anti-SARS-CoV-2 activity.

## MATERIALS & METHODS

### Chemicals and inhibitors

(+)-JQ-1 (Cat no. SML1524-5MG) was purchased from Sigma. ABBV-075 (Cat no. S8400) (ClinicalTrails.gov Identifier: <u>NCT02391480</u>), OTX015 (Cat no. S7360) (ClinicalTrails.gov Identifier: <u>NCT02698176</u>), ARV-825 (Cat no. S8297) and ML385 (Cat no. S8790) were purchased from Selleckchem. Dimethyl sulfoxide (DMSO) (Item no. 10127403) was purchased from Thermo Fisher Scientific.

### Cell lines

Calu-3 (ATCC HTB-55), CaCo-2 (ATCC HTB-37), Vero E6 (ATCC CRL-1586) and parental HEK293T cells (ATCC CRL-3216) were cultured in Dulbecco’s Modified Eagle’s Medium (DMEM) supplemented with 10% heat-inactivated fetal calf serum (FCS), 100 U/ml Penicillin-Streptomycin and 2mM L-glutamine (Gibco, UK) (hereafter referred to as 10% DMEM) at 37°C/5% CO_2_. Primary bronchial airway epithelial cells were cultured in air-liquid interface (ALI) mode in the provided ALI medium at 37 °C/5% CO_2_. Unless stated otherwise, all cell cultures were maintained at 37°C/5% CO_2_.

### Human bronchial airway epithelial cells (Air-liquid interface cultures)

Human bronchial airway epithelial cells (hBAECs) were isolated from explanted lungs obtained from the Hannover Lung Transplant Program after patients’ informed consent, ethical vote 2923-2015. For isolation of hBAECs, human bronchial tissue was cut into small pieces in Hank’s buffer (Thermo Fisher Scientific) containing 0.18% protease XIV and incubated for two hours at 37°C. After thorough pipetting with a 25/50 ml serological pipette, cell solution was filtered through a 100 μm cell strainer (Corning) to remove clumps and 10 ml RPMI supplemented with 10% FCS (Thermo Fisher Scientific) was added. After centrifugation for 10 min at 500*g* and 4°C, supernatant was removed and cells were resuspended in SAEGM™ (PromoCell) + Primocin (InvivoGen) + Penicillin-Streptomycin (P/S) (Sigma-Aldrich). For ALI cultures, 200,000 hBAECs were seeded onto PureCol- (Advanced BioMatrix) coated 12- well inserts (Greiner Bio-One) in SAEGM™ + Primocin + P/S. 48 hours post seeding, culture medium in apical and basal chambers was changed to PneumaCult-ALI medium (STEMCELL Technologies). Air lift was performed 48 hours later by gently removing medium from the apical chamber. Homogenous distributed cilia were visible three weeks after air lift and inserts were used for infections.

### Cell viability assays

Calu-3 cells (6×10^5^ cells/ml) seeded overnight in 96-well plates were treated with serial dilutions (prepared in culture medium) of the iBETs or corresponding DMSO controls for 72 hours; with PBS wash, medium change and fresh drug administration after every 24 hours. Post-treatment, cells were subjected to viability assays using the CellTiter-Glo Luminescent Cell Viability Assay Kit (Promega, Germany) for Calu-3 cells and CellTiter-Glo 3-D Cell Viability Assay Kit (Promega, Germany) for hBAECs according to the manufacturer’s protocol. The raw data from the test samples were background subtracted, normalised to naïve cells, and analysed using GraphPad Prism v9 (LaJolla, CA, USA) as previously described (Mhlekude et al., 2021).

### Virus production

SARS-CoV-2 (passage (P) 1, BetaCoV/Munich/BavPat1/2020|EPI_ISL_406862) (Wölfel et al., 2020), SARS-CoV (P1, HKU-39849 Hong Kong) (Lau et al., 2005; Zeng et al., 2003), and MERS-CoV (P2, EMC/2012) (Zaki et al., 2012) stocks were propagated in Vero E6 cells to generate P2, P2, and P3 stocks respectively. The virus stock production and quality control by Next Generation Sequencing were conducted as previously described (Niemeyer et al., 2021). All infectious pathogens were handled under biosafety level three (BSL-3) conditions with respiratory personal protection equipment.

### Virus infection of Calu-3 cells

Calu-3 cells (6×10^5^ cells/ml) seeded overnight in 12-well plates were treated with serial dilutions of iBETs or corresponding DMSO controls for 48 hours; with PBS wash, medium change and fresh drug administration every 24 hours. Post-treatment, cells were inoculated with viruses (SARS-CoV and SARS-CoV-2, MOI 0.1; and MERS-CoV, MOI 0.0005) and incubated for one hour. After infection, cells were washed with PBS, supplied with fresh medium and iBETs and incubated for 24 hours. The next day, 50 μl of supernatant was collected into 300 μl of RAV1 lysis buffer (Machery-Nagel, Germany) for viral RNA extraction and 100 μl into 100 μl of 0.5% gelatine medium for plaque assays. Cells were washed with PBS, trypsinized and reconstituted in culture medium, from which 50 μl was collected into 300 μl of RAV1 lysis buffer (Machery-Nagel, Germany) to isolate cell-associated RNA. RNA samples were stored at −20°C and plaque assay samples at −80°C until sample processing.

### Virus infection of human bronchial airway epithelial cells (hBAECs)

JQ-1 or DMSO was added into the culture medium (PneumaCult-ALI) in the basal compartment of cells cultured in transwells and incubated for 48 hours, followed by medium exchange and new drug administration after 24 hours as mentioned above. Post-treatment, cells were washed three times with pre-warmed PBS to remove mucus from the apical compartment and inoculated with SARS-CoV-2 (2×10^4^ PFU diluted in OptiPRO medium), followed by incubation for 90 minutes. The inoculum was then aspirated and cells washed with PBS three times. The transwells were then transferred into new plates with the drug-containing culture medium. To harvest samples for zero time point, 200 μl of culture medium was immediately added onto the cells in the apical compartment and incubated for 15 min to elute the virus particles. Following incubation, 50 μl of virus eluate was collected into 300 μl RAV1 lysis buffer (Machery-Nagel, Germany) for viral RNA extraction and 100 μl into 100 μl of 0.5% gelatine medium for plaque assays, after which the eluate residues were aspirated from the apical compartment and followed by incubation of cells. This harvesting procedure was repeated every 24 hours for kinetic experiments. The samples were stored at −20°C and −80°C, respectively, until processing.

### Quantitative real-time PCR

Viral RNA from the supernatant was extracted using the NucleoSpin RNA Virus Isolation Kit (Macherey-Nagel, Germany) according to the manufacturer’s protocol. To quantify viral RNA copies, a 12.5 μl reaction/well was prepared in 4titude® 96-Well Semi-Skirted PCR Plates for LC480 (Roche) using a SuperScript III One-Step RT-PCR Kit (Invitrogen, Germany). Reverse transcription of the viral RNA and amplification of the cDNA was conducted in a LightCycler 480 System (Roche, Germany) using the following protocol: 55°C, 10 min; 95°C, 3 min; 95°C, 15 sec; 58°C, 30 sec and 40°C, 30 sec for 45 cycles and qPCR primers against SARS-CoV and SARS-CoV-2 E gene and probes (Corman et al., 2020). The absolute quantification of the viral RNA copies was calculated by a standard curve method using viral RNA standards (Corman et al., 2020).

### Plaque titration assays

Seeded Vero E6 cells (4 × 10^5^ cell/ml) were left overnight in 24-well plates. The virus-containing samples (50 μl) were serially diluted in 450 μl aliquots of serum-free Opti-Pro medium (Gibco, UK), after which 200 μl from each dilution were added in duplicates to the cells, followed by one hour incubation. Post-infection, viral inoculum was aspirated from the cells, cells washed with PBS, and overlaid with 2.4% Avicel (FMC BioPolymers, Germany) mixed with 2xDMEM (1:1 ratio), followed by incubation for 72 hours. The overlay was aspirated from the cells, after which they were fixed for 30 min with 6% formaldehyde and stained with crystal violet solution (0.2% crystal violet, 2% ethanol and 10% formaldehyde) for 20 min. Plaque Forming Units (PFU) were determined from at least two dilutions for which distinct plaques were detectable and expressed as PFU/ml.

### Flow cytometry

JQ-1 and DMSO-treated Calu-3 cells were infected with GFP-tagged SARS-CoV-2 (synSARS-CoV-2-GFP-P2A-ORF7a clone #41, (Thi Nhu Thao et al., 2020) (MOI 0.25) following the above-mentioned protocol. Subsequently, cells were trypsinized, PBS-washed and fixed for 90 minutes with 4% PFA. GFP signal was quantified by flow cytometry in FACSCelesta (BD Bioscience, Germany) and analysed by FlowJo v10.8 (Tree Star, Ashland, Oregon, USA).

### Immunoblotting

Calu-3 cells were lysed in 60 μl RIPA Lysis Buffer (Thermo Fisher Scientific, Germany) supplied with 1 % Protease Inhibitor Cocktail Set III (Merck Chemicals, Germany) for 30 min at 4 °C. Subsequently, cell lysates were pelleted and protein concentration in supernatant was determined by BCA protein assay (ThermoFisher Scientific). 20 μg total protein from each sample was mixed with 4x Laemmli buffer, which was supplemented with 10% beta-mercaptoethanol, then boiled for ten minutes at 95°C to ensure protein denaturation. Proteins were resolved on 6% SDS-PAGE gel and transferred to a nitrocellulose membrane (0.45 μm pore size, GE Healthcare) by Trans-Blot Turbo system (BioRad). Membranes were blocked with 5% dried milk in 0.1% PBS-Tween (0.9% NaCl, 10 mM Tris-HCl [pH 7.5], 0.1% Tween 20) for 30 min at room temperature. Blocked membranes were incubated with the primary antibodies against ACE2 (#AF933, R&D Systems) and beta-actin (#A5316, Sigma-Aldrich). Secondary antibodies conjugated with horseradish peroxidase (HRP) were used for chemiluminescence-based detection by Fusion Fx7 (Peqlab Biotechnologie GmbH). Detection was performed using SuperSignal™ West Femto substrate (ThermoFisher Scientific).

### Production of pseudotyped lentivirus particles and transduction assays

SARS-CoV-2-S- and VSV-G-pseudotyped lentiviral particles (Hu et al., 2020) were produced by calcium phosphate-based transfection of HEK293T cells with the packaging plasmid pCMV ΔR8.91 (Zufferey et al., 1997), the lentiviral transfer plasmid pCSII-EF-luciferase (Agarwal et al., 2006), and pCMV-VSV-G (Stewart et al., 2003) using the CalPhos Mammalian Transfection Kit (Takara Bio Company, Germany) according to the manufacturer’s protocol. Dose-dependent treatment of Calu-3 cells with JQ-1, followed by transduction with pseudovirus particles in 96-well plates was conducted under BSL-2 conditions, after which the infection efficiency was analysed luminometrically in Synergy™ HTX Multi-Mode Microplate Reader (BioTek Instruments, Inc.) using the Luciferase Assay System (Promega, Germany) according to the manufacturer’s protocol.

### ATAC-seq

Bulk ATAC-seq and RNA-seq were conducted in parallel from the identical infection experiment. Calu-3 cells were treated with JQ-1 and DMSO, respectively, for 24 hours, infected with SARS-CoV-2 (MOI 0.1) under continuous treatment for another 24 hours. Uninfected but treated, and naïve Calu-3 cells were used as reference. Post-infection, ATAC-seq libraries were prepared from 50,000 cells per replicate using Illumina Tagment DNA Enzyme and Buffer Kit (Illumina 20034197) according to the Omi-ATAC-seq protocol (Corces et al., 2017) with the following minor optimizations. Briefly, 5 μl from the partially amplified barcoded fragments was subjected to SYBR Green qPCR in a LightCycler 480 System (Roche, Germany) using the FastStart Essential DNA Green Master Mix (Roche, Germany) according to the manufacturer’s protocol. Amplification was conducted for 20 cycles using a universal forward primer and the sample-specific barcoded reverse primers (Buenrostro et al., 2013). Amplification curves from SYBR Green qPCR were generated and used to determine the number of cycles that give ⅓ of the maximum fluorescence. Final library preparation was conducted by conventional PCR for 8-12 cycles. The amplified libraries were subjected to a single left-sided bead purification using the AMPure XP magnetic beads (Beckman Coulter, Germany). Libraries were sequenced on SP lane of NovaSeq 6000 System (Illumina) at the MDC/BIH Genomic Core Facility to generate 40 million 75-nucleotide paired-end reads per sample.

### RNA-seq

Total RNA was isolated from 3×10^5^ cells per replicate using Direct-Zol RNA Miniprep Kit (Zymo Research, USA) according to the manufacturer’s protocol and shipped in dry ice to the MDC/BIH Genomic Core Facility for quality assessment using TapeStation (Agilent Technologies). Libraries were prepared using TrueSeq Stranded mRNA kit to generate Illumina-compatible libraries by following the manufacturer’s protocol (Illumina). Libraries were sequenced on SP lane of NovaSeq 6000 System (Illumina) to generate 40 million 75-nucleotide paired-end reads per sample.

### Liquid chromatography tandem mass spectrometry (LC-MS/MS)

Treatment of Calu-3 cells and infection with SARS-CoV-2 were performed following the above-mentioned protocol. Treated but uninfected and naïve cells served as references. Preparation of cell lysate samples and protein quantification were conducted as mentioned above, from which 50 μg of proteins per sample were prepared and resolved on 7.5% linear SDS-PAGE gels as described elsewhere (Mhlekude et al., 2021). The in-gel protein digestion and liquid chromatography coupled with tandem mass spectrometry (LC-MS/MS) experiments were performed as previously described (Jochim et al., 2011).

### Bioinformatics analysis

For RNA-seq, reads were aligned to the human genome version GRCh38 and counted using STAR (Dobin et al., 2013). Differential expression analysis was conducted using DESeq2 version 1.30 (Love et al., 2014). Transcription factor analysis was performed using the R package Dorothea (Garcia-Alonso et al., 2019) For ATAC-seq, reads were aligned to the human genome GRCh38, after which the peaks were called using MACS2 v. 2.2.7.1 (Zhang et al., 2008). For differential TF binding analysis, the R package DiffBind v. 3.0.15 (Stark and Brown) was used. Motif search analysis was performed using the MEME suite (Bailey et al., 2015) and DREME (https://meme-suite.org/meme/doc/dreme.html). Gene set enrichment analysis was performed with the R package cluster Profiler v. 3.18 (Yu et al., 2012). All results were corrected for multiple testing using the Benjamini-Hochberg procedure (Benjamini and Hochberg, 1995). Genes were annotated as involved in IFN signalling, the cell cycle, or autophagy by referring to the Reactome Interferon Signalling geneset (R-HSA-913531), the Reactome Cell Cycle geneset (R-HSA-1640170), and the KEGG Autophagy - animal geneset (hsa04140), respectively. Genes identified as targets for NRF2 signalling were identified based on a previous report (Olagnier et al., 2020). LC-MS/MS data analysis was conducted as previously described (Mhlekude et al., 2021).

### Statistical analysis

If not stated otherwise, bars show the arithmetic mean of indicated amount of repetitions. Error bars indicate standard error of the mean (SEM) from the indicated amount of individual experiments. Statistical significance was calculated by performing a Student’s t-test using GraphPad Prism. *P* values ≤0.05 were considered significant: □0.05 (*), □0.0021 (**), □0.0002 (***), □0.0001 (****) and n.s. = not significant (≥0.05).

## Data and code availability

All code used to perform the analysis of the RNAseq and ATACseq data will be available at https://github.com/GoffinetLab/SARS-CoV-2_JQ1-Antiviral-Study. Raw and processed OMICs data will be deposited on the GEO database.

## RESULTS

### Prophylactic administration of inhibitors of bromodomain and extra-terminal proteins (iBET) inhibit SARS-CoV-2 infection in lung epithelial Calu-3 cells

Inhibition of SARS-CoV-2 infection by several iBET compounds has been reported in different infection models (Gilham et al., 2021; Mills et al., 2021; Samelson et al., 2022). To gain more insights into their antiviral potency in the context of SARS-CoV-2 infection, we compared antiviral activity of four iBETs, three (JQ-1, ABBV-075, and OTX-015) of which compete with the acetylated molecules for binding to the bromodomains of the BET proteins, and one (ARV-825) that targets BET proteins (dBET) carrying proteolysis-targeting chimera (PROTAC) sequences to proteasomal degradation. iBET treatment of Calu-3 cells prior to SARS-CoV-2 infection led to a dose-dependent reduction of viral genomic RNA quantities (Fig. 1A and Sup. Fig. 1C) and infectious titers (Fig. 1B and Sup. Fig. 1D) in the supernatant in the absence of detectable cell toxicity (Sup. Figs. 1A and 1E). Of the investigated iBET candidates, JQ-1 (IC_50_ = 0.290 μM) and ABBV-075 (IC_50_=0.132 μM) were more potent than OTX-015 (IC_50_ = 3.553 μM) and ARV-825 (IC50 = 1.431 μM) against SARS-CoV-2 infection in Calu-3 cells. Prophylactic administration of JQ-1 to hBAECs reduced SARS-CoV-2 viral RNA quantities by 24.8-fold (Fig. 1C) and infectious titers by 28.2-fold (Fig. 1D) without detectable toxicity (Sup. Fig. 1B), demonstrating the relevance of iBET-mediated anti-SARS-CoV-2 activity in a physiologically relevant primary cell model (Samelson et al., 2022). JQ-1 treatment rendered Calu-3 cells 2.7-fold less susceptible to infection by recombinant GFP-expressing SARS-CoV-2, reflecting the consistency of JQ-1-mediated anti-SARS-CoV-2 activity across different readouts (Fig. 1E-F).

**Figure 1.**
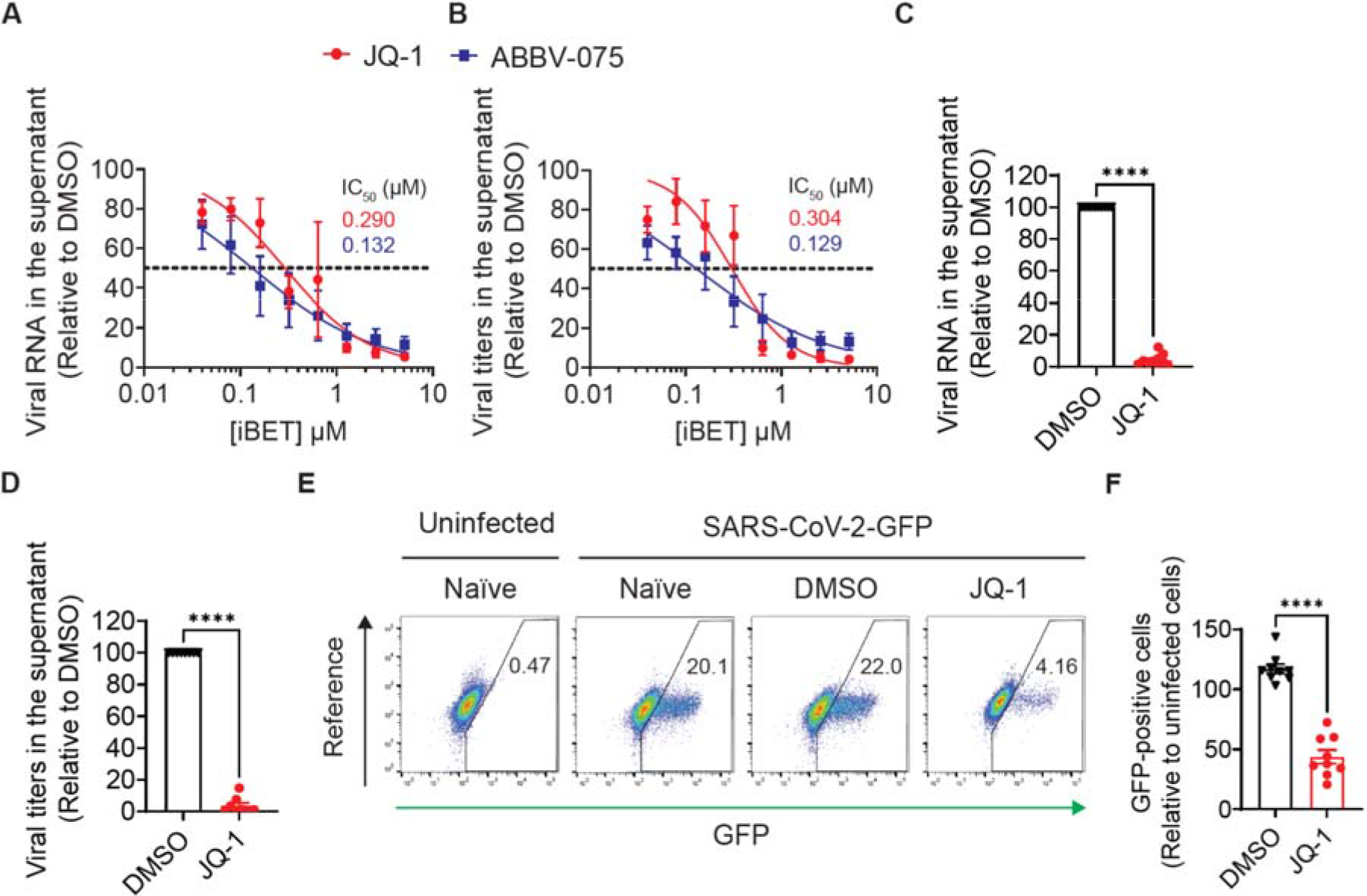
Prophylactic administration of inhibitors of bromodomain and extra-terminal proteins (iBET) inhibit SARS-CoV-2 infection in lung epithelial Calu-3 cells. (**A**-**B**) Calu-3 cells were pretreated with indicated concentrations of iBETs for 48 hours, followed by infection with SARS-CoV-2 (MOI 0.1) under continuous presence of the drug. At 24 hours post infection, relative (**A**) SARS-CoV-2 RNA and (**B**) infectious titers in supernatants were quantified from three independent experiments. (**C**-**D**) hBAECs were pretreated with JQ-1 (2.56 μM) for 48 hours, followed by infection with SARS-CoV-2 (2×10^4^ PFUs) under continuous presence of the drug. At 48 hours post-infection, (**C**) abundance of viral RNA from duplicates of five independent experiments and (**D**) infectivity from duplicates of four independent experiments were quantified in supernatants. Data shown are relative to DMSO treatment. (**E**-**F**) Calu-3 cells were pretreated with JQ-1 for 48 hours prior to infection with a recombinant, GFP-expressing SARS-CoV-2 (MOI 0.25). Shown are (**E**) representative FACS dot plots and (**F**) quantification of percentage of GFP-positive cells relative to DMSO treatment at 24 hours post infection, from three independent experiments with triplicates each. Unpaired nonparametric Student *t*-test was used to compare the means between the experimental groups. Error bars represent the arithmetic mean±SEM.

### JQ-1 exhibits a subgenera-specific antiviral activity among *Betacoronaviridae* in lung epithelial Calu-3 cells

To investigate whether the iBET-mediated anti-SARS-CoV-2 activity extends beyond SARS-CoV-2 parental strains, which have been the main focus of the previous studies (Gilham et al., 2021; Mills et al., 2021; Samelson et al., 2022), we compared the antiviral potency of JQ-1 in Calu-3 cells infected with a panel of ß-coronaviruses. JQ-1 potently and dose-dependently inhibited infection by SARS-CoV, SARS-CoV-2_B.1_, SARS-CoV-2_B.1.1.7_ and SARS-CoV-2_BA.2_, in contrast to MERS-CoV, as indicated by the reduction of viral RNA copies (Fig. 2A) and infectious titers (Fig. 2B) in culture supernatants. These data suggest that JQ-1 exhibits a subgenera-specific antiviral activity, which is directed towards infection by *Sarbecoviruses* and not *Merbecoviruses* to which MERS-CoV belongs.

**Figure 2.**
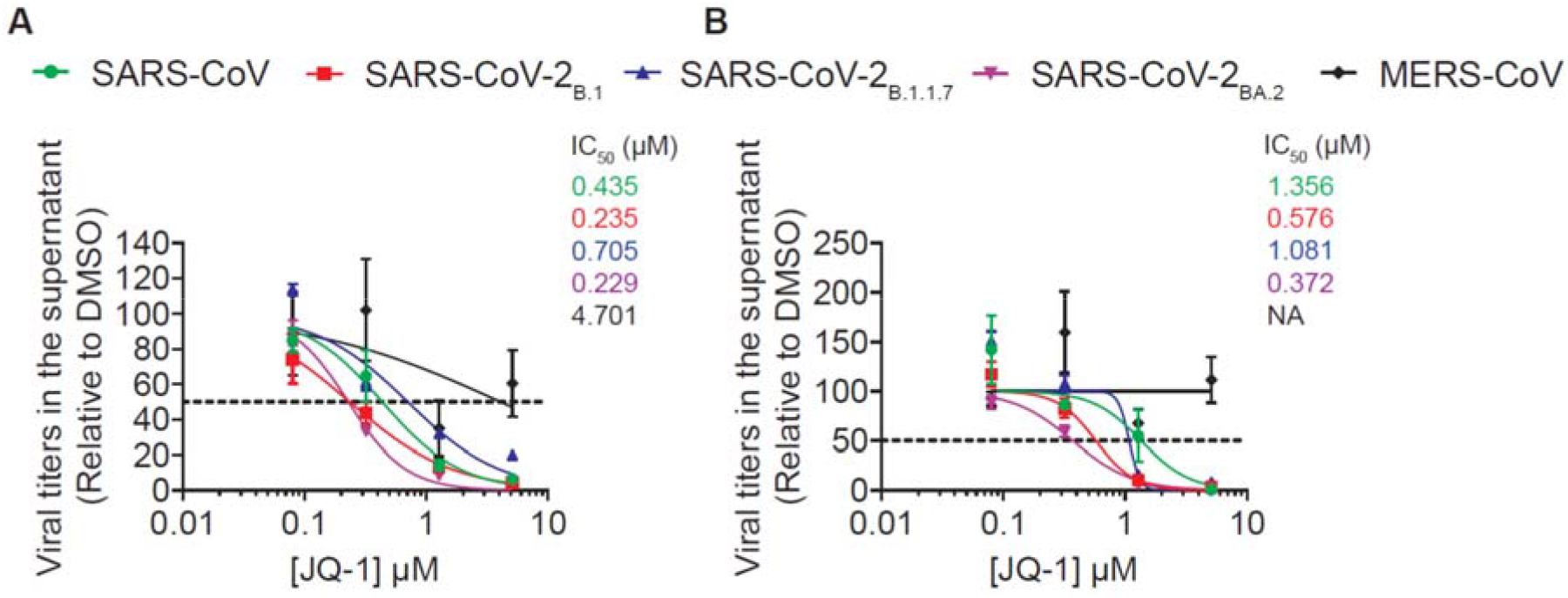
JQ-1 exhibits a subgenera-specific activity among *Betacoronaviridae* in lung epithelial Calu-3 cells. Calu-3 cells were pretreated with indicated concentrations of JQ-1 prior to infection with indicated viruses. Shown are DMSO-normalised (**A**) viral RNA abundance and (**B**) infectivity in supernatants harvested at 24 h.p.i. with *Sarbecoviruses* (MOI 0.1) and *Merbecovirus* (MOI 0.0001). Error bars represent the mean±SEM.

### JQ-1 exhibits a cell-directed anti-SARS-CoV-2 activity in lung epithelial Calu-3 cells

Next, we compared the effect of JQ-1-mediated anti-SARS-CoV-2 activity when administered prophylactically and at the time point of infection. Focusing on a single round of infection, we found that JQ-1 inhibited SARS-CoV-2 replication when administered prophylactically in Calu-3 cells, while it failed to inhibit infection when administered to cells during infection (Fig. 3A). PCR-based quantification of viral RNA copies showed that JQ-1-mediated reduction of viral RNA copies in the supernatant of infected Calu-3 cells is accompanied by, and probably a consequence of, reduced quantities of cell-associated viral RNA (Fig. 3B). The ratio of extracellular to total viral RNA indicated that JQ-1 impaired release of SARS-CoV-2 virions (Fig. 3C), without *per se* reducing the specific particle infectivity of secreted particles (Fig. 3D). In accordance with previous reports (Gilham et al., 2021; Mills et al., 2021; Samelson et al., 2022), JQ-1 treatment reduced steady-state levels of ACE2 expression (Fig. 3E), an observation that was accompanied by reduced susceptibility of Calu-3 cells to transduction with SARS-CoV-2 spike, but not VSV-G-pseudotyped lentiviral particles (Sup. Fig. 3F). Together, these data suggest that JQ-1 exhibits a cell-directed anti-SARS-CoV-2 activity involving, but likely not restricted to, downregulation of ACE2 expression.

**Figure 3.**
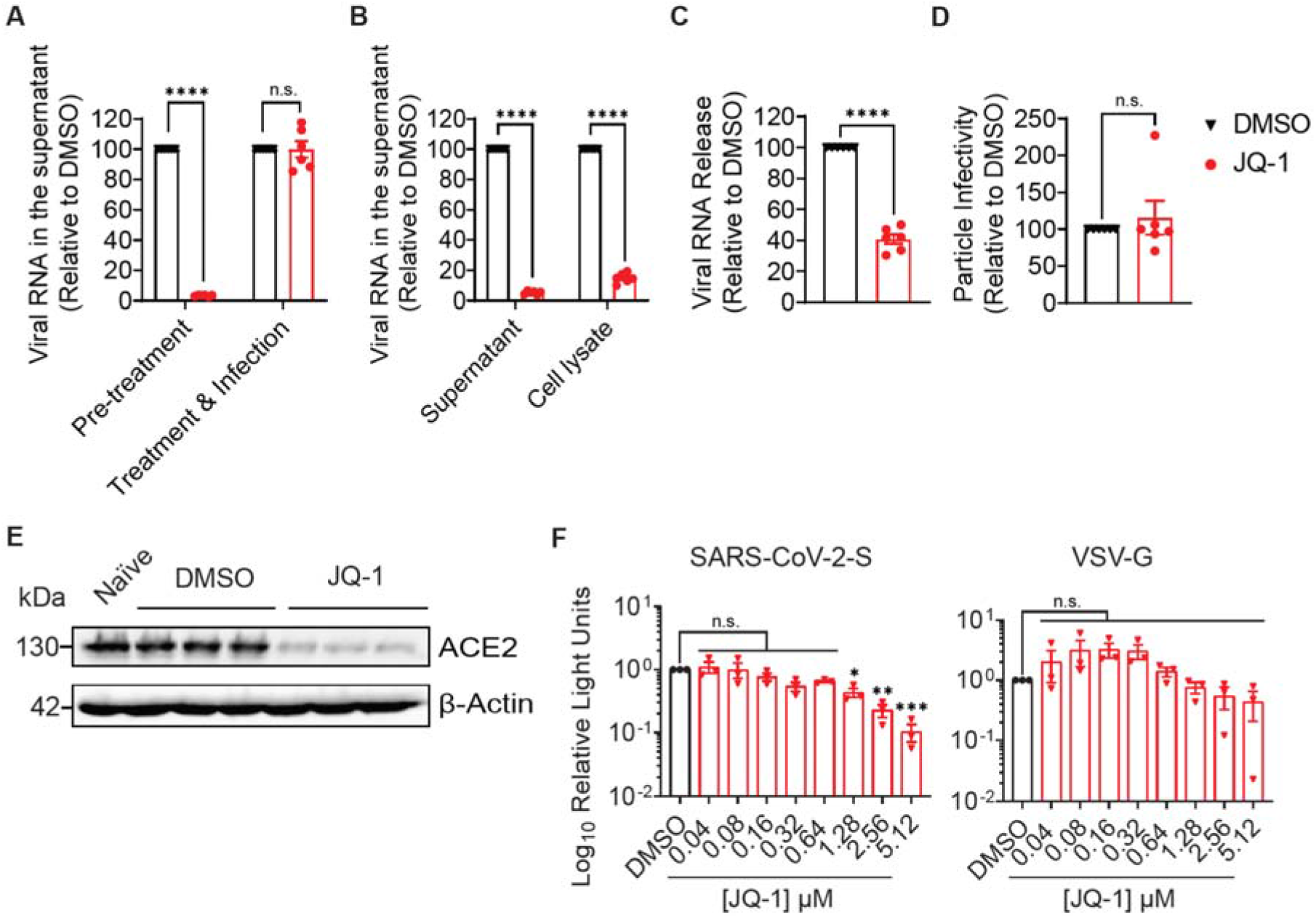
JQ-1 exhibits a cell-directed anti-SARS-CoV-2 activity in lung epithelial Calu-3 cells. (**A**) Calu-3 cells were either pre-treated with JQ-1 (2.56 μM) for 48 hours, followed by SARS-CoV-2 infection (MOI 0.1) and continued treatment (Pre-treatment), or treatment was initiated at the time point of infection (Treatment & Infection). 24 hours post-infection, relative viral RNA abundance in supernatants was quantified. (**B-D**) The relative efficiency of different steps of the replication cycle were quantified following JQ-1 treatment and infection of Calu-3 cells, including (**B**) viral RNA abundance in supernatants and cell lysates, (**C**) particle release and (**D**) specific particle infectivity. The error bars represent the arithmetic mean±SEM from duplicates of three independent experiments. Viral release was calculated by dividing the concentration of viral RNA in the supernatant by the sum of the concentrations of the viral RNA in the supernatant and cell lysate. Particle infectivity was calculated by dividing the viral titers by the corresponding viral RNA concentration in the supernatant. **(E)** Immunoblot analysis of ACE2 expression in JQ-1-treated Calu-3 cells **(F)** Luminometric quantification of transduction efficiency of JQ-1-treated Calu-3 cells using lentiviral pseudoparticles decorated with SARS-CoV-2-S and VSV-G proteins. Error bars represent the mean±SEM from three independent experiments. Unpaired nonparametric Student *t*-test was used to compare the means between the experimental groups.

### SARS-CoV-2 infection and JQ-1 treatment modulate the chromatin regulatory landscape in lung epithelial Calu-3 cells

To investigate the chromatin regulatory landscape that associates with JQ-1-mediated SARS-CoV-2 inhibition, we subjected Calu-3 cells to bulk ATAC-seq. Principal component analysis (PCA) showed clustering of samples according to their experimental groups, reflecting comparable chromatin profiles between individual samples in each group (Sup. Fig. 2A). JQ-1 treatment imposed larger changes than SARS-CoV-2 infection. Samples from uninfected DMSO-treated cells closely clustered with samples from naïve cells, suggesting that DMSO treatment induced minimal changes to the host chromatin profile. Compared to the control groups, JQ-1 administration to Calu-3 cells increased accessibility to the transcriptional start sites (TSS) irrespective of the infection status, followed by SARS-CoV-2 infection, as indicated by the increase in the density of peaks mapping to the TSSs (Fig. 4A), illustrating the differential ability of JQ-1 treatment and SARS-CoV-2 infection to remodel chromatin accessibility to the TSSs. Annotation of the accessible peaks to the genomic features in the chromatin showed a large coverage for promoter sequences located within one kb from the TSS regions across all experimental groups (Fig. 4B). However, irrespective of the infection status, JQ-1 treatment shifted a proportion of the accessible regions away from the promoter sequences located within one kb from the TSS regions to the introns and distal intergenic regions. This suggests that regulation of transcription under JQ-1 treatment is driven by a more evenly-distributed accessibility of genomic features compared to SARS-CoV-2 infection or absence of JQ-1, where transcriptional regulation is predominantly driven by the proximal regulatory cis-elements in the promoters. Hierarchical clustering of significantly regulated peaks in each experimental group revealed distinct chromatin accessibility profiles, modulated distinctly by either SARS-CoV-2 infection or JQ-1 treatment, with some commonly upregulated peaks resulting from both SARS-CoV-2 infection and JQ-1 treatment, respectively (Fig. 4C).

**Figure 4.**
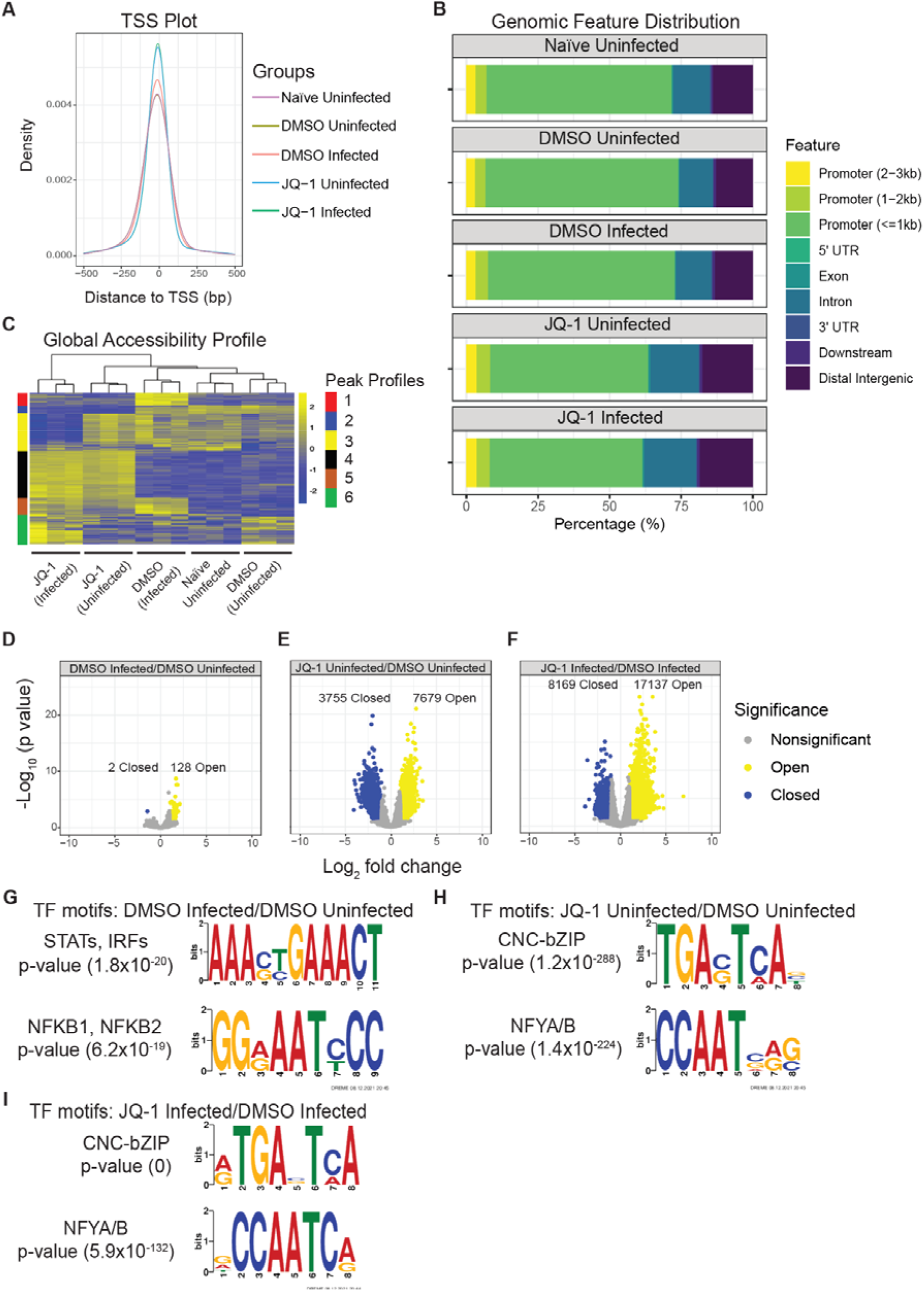
SARS-CoV-2 infection and JQ-1 treatment modulate the chromatin regulatory landscape in lung epithelial Calu-3 cells. **(A)** TSS plot depicting the density and distribution of accessible ATAC-seq peaks around transcription start sites within a window of −500 to 500 bp. **(B)** Analysis of the genomic features annotated to accessible ATAC-seq peaks in the chromatin. (**C**) Heatmap hierarchical clustering of differentially accessible ATAC-seq peaks scaled as z-score across rows. Volcano plots showing relative log_2_FC and statistical significance of differentially regulated ATAC-seq peaks in (**D**) DMSO-treated infected versus DMSO-treated uninfected, (**E**) JQ-1-treated uninfected versus DMSO-treated, uninfected and (**F**) JQ-1-treated infected versus DMSO-treated, infected contrasts. Significantly (FDR of ≤0.05) regulated peaks are indicated. TF motif enrichment analysis from the accessible ATAC-seq peaks in (**G**) DMSO-treated, infected versus DMSO-treated uninfected, (**H**) JQ-1-treated uninfected versus DMSO-treated, uninfected and (**I**) JQ-1-treated infected versus DMSO-treated, infected contrasts. The TF motifs were identified using the DREME algorithm, where the height of the letters represents the frequency of each base in the motif.

Volcano plots of the ATAC-seq data revealed that SARS-CoV-2 infection only mildly modulated the chromatin accessibility profile (significant changes in 130 ATAC-seq peaks, 128 of which were accessible while only 2 were inaccessible) (Fig. 4D). In contrast, JQ-1 treatment drastically altered the chromatin accessibility landscape (Fig. 4E, significant changes of 11,434 ATAC-seq peaks, 7679 of which were accessible while 3,755 were inaccessible). Strikingly, combination of JQ-1 treatment and infection induced 2.2-fold increased change to the accessibility of genomic regions when compared to JQ-1 treatment only (Fig. 4F, significant changes in 25,306 ATAC-seq peaks, whereby 17,137 were accessible while 8,169 were inaccessible), implying that changes induced by JQ-1 pretreatment are quantitatively superior to those induced by infection.

As expected (Wilk et al., 2021; You et al., 2021), compared to uninfected cells, pathways that drive innate immune responses were enriched in the accessible ATAC-seq peaks from SARS-CoV-2-infected cells (Sup. Fig 2B). Accessible ATAC-seq peaks from JQ-1-treated cells showed no significant association with any particular biological pathway when compared to DMSO-treated cells (Sup. Fig 2B-C), suggesting that JQ-1 induced a rather global change rather than alterations of specific biological processes. On the other hand, when compared to infected cells, accessible ATAC-seq peaks from infected cells in the presence of JQ-1 were mostly enriched with ribosome biogenesis and RNA processing pathways (Sup. Fig 2B), including RNA splicing reported to be inhibited by SARS-CoV-2 (Banerjee et al., 2020). Of note, nonsense-mediated mRNA decay, a cellular RNA surveillance pathway that exhibits a broad antiviral activity (Balistreri et al., 2014; Wada et al., 2018), was among the highly enriched pathways in the accessible peaks (Sup. Fig 2B). Conversely, pathways associated with sensory perception of smell were downregulated in the accessible peaks from SARS-CoV-2-infected cells, both in the presence of DMSO and JQ-1 (Sup. Fig 2C), but not by JQ-1 *per se*, suggesting an inability of JQ-1 to hamper SARS-CoV-2-mediated downregulation of the smell receptor signalling pathway. Furthermore, accessible peaks from JQ-1-treated infected cells were associated with downregulation of the neuropeptide signalling pathway (Sup. Fig 2C), as opposed to infection or JQ-1 treatment, suggesting JQ-1’s potential to interfere with neuronal communication specifically in the infection context. Together, these data show that SARS-CoV-2- and JQ-1-mediated modulation of the chromatin accessibility landscape occurs at multiple levels, with differing magnitudes and breadth.

We next searched for transcription factor (TF) binding motifs that were significantly enriched in the accessible ATAC-seq peaks using the DREME algorithm, which is designed to find short and multiple nonredundant binding motifs of eukaryotic TFs and calculate their statistical significance (Bailey, 2011). In infected cells, accessible peaks were enriched for binding motifs for inflammatory TF families, including Signal Transducer and Activator of Transcription (STAT), Interferon Regulatory Factors (IRF) and Nuclear Factor-kappa B (NFkB), which drive IFN signalling and induction of pro-inflammatory cytokines, respectively (Park and Iwasaki, 2020) (Fig. 4G and Sup. Table 1A), suggesting increased accessibility of STAT-, IRF- and NFκB-binding sites to induce IFN response pathways and production of pro-inflammatory cytokines (Bülow Anderberg et al., 2021; Ramasamy and Subbian, 2021). Unlike in SARS-CoV-2 infection, the accessible peaks in JQ-1-treated cells showed significant enrichment of several binding motifs of a wide variety of TF families, suggesting a broader modulation of gene expression by JQ-1 over SARS-CoV-2 infection (Sup. Table 1B-C). Independent of infection, among many others, peaks from JQ-1-treated cells were enriched with binding motifs for TF families such as CNC-bZIP and NF-Y (Fig. 4H-I, Sup. Table 1B- C), which induce antiviral cellular cytoprotective responses (He et al., 2020; Olagnier et al., 2020) and more accessible chromatin (Oldfield et al., 2014, 2019), respectively. Detection of these enrichments, independent of SARS-CoV-2 infection, argues for a dominant role of JQ-1 treatment on transcription modulation via the CNC-bZIP and NFY TF families. Secondly, motifs annotated to a superfamily of steroid-induced nuclear receptor (NR) TFs, whose signalling drives cellular RNA splicing processes (Auboeuf et al., 2004; Elhasnaoui et al., 2021), were also enriched in JQ-1-treated groups irrespective of the infection status (Sup. Table 1B-C). However, JQ-1 treatment in the presence of SARS-CoV-2 infection (Sup. Table 1C) displayed significant enrichment with TF binding motifs for a wide range of TF families compared to the absence of infection (Sup. Table 1B), suggesting that SARS-CoV-2 infection influences JQ-1-mediated TF profile and likely fine-tunes the JQ-1-mediated transcriptome. Interestingly, compared to uninfected cells, peaks from uninfected JQ-1-treated cells were significantly enriched with a binding motif for NFκB-1/NFκB-2 TFs (Sup. Table 1B), despite JQ-1 being an inhibitor of the NFκB-1-mediated canonical pathway that induces cytokine production (Samelson et al., 2022). Together, these data illustrate the genome-wide TF binding profiles that drive the transcriptional programs governing SARS-CoV-2 infection- and JQ-1-mediated changes.

### SARS-CoV-2- and JQ-1-mediated modulations of the chromatin accessibility landscape underlie the transcriptomic and proteomic changes in lung epithelial Calu-3 cells

Next, we subjected the samples to bulk RNA-seq and mass spectrometry to establish their transcriptomic and proteomic profiles, respectively, including the downstream effects of SARS-CoV-2- and JQ-1-mediated modulations on the chromatin accessibility landscape. PCA of the RNA-seq data revealed that the samples clustered according to their experimental groups, reflecting similar transcriptomic profiles between replicates in each group (Sup. Fig. 3A). As in ATAC-seq (Sup. Fig. 2A), JQ-1 treatment induced more important changes than SARS-CoV-2 infection. Samples from naïve and uninfected DMSO-treated cells clustered close to each other, with minimal effect of DMSO treatment on the cellular transcriptome (Sup. Fig. 3A).

SARS-CoV-2 infection induced significant changes to the expression of 6866 genes, of which 3323 were upregulated and 3543 downregulated (Fig. 5A). In the absence of infection, JQ-1 treatment induced significant changes to the expression of 10,778 genes, of which 5285 were upregulated and 5493 were downregulated (Fig. 5B). On the other hand, when compared to infection only, JQ-1 treatment of infected cells induced significant changes to the expression of 11,195 genes, with 5128 upregulated and 6067 downregulated (Fig. 5C). Again, these data suggest that JQ-1 treatment modulates the host transcriptome to a larger extent compared to SARS-CoV-2 infection.

**Figure 5.**
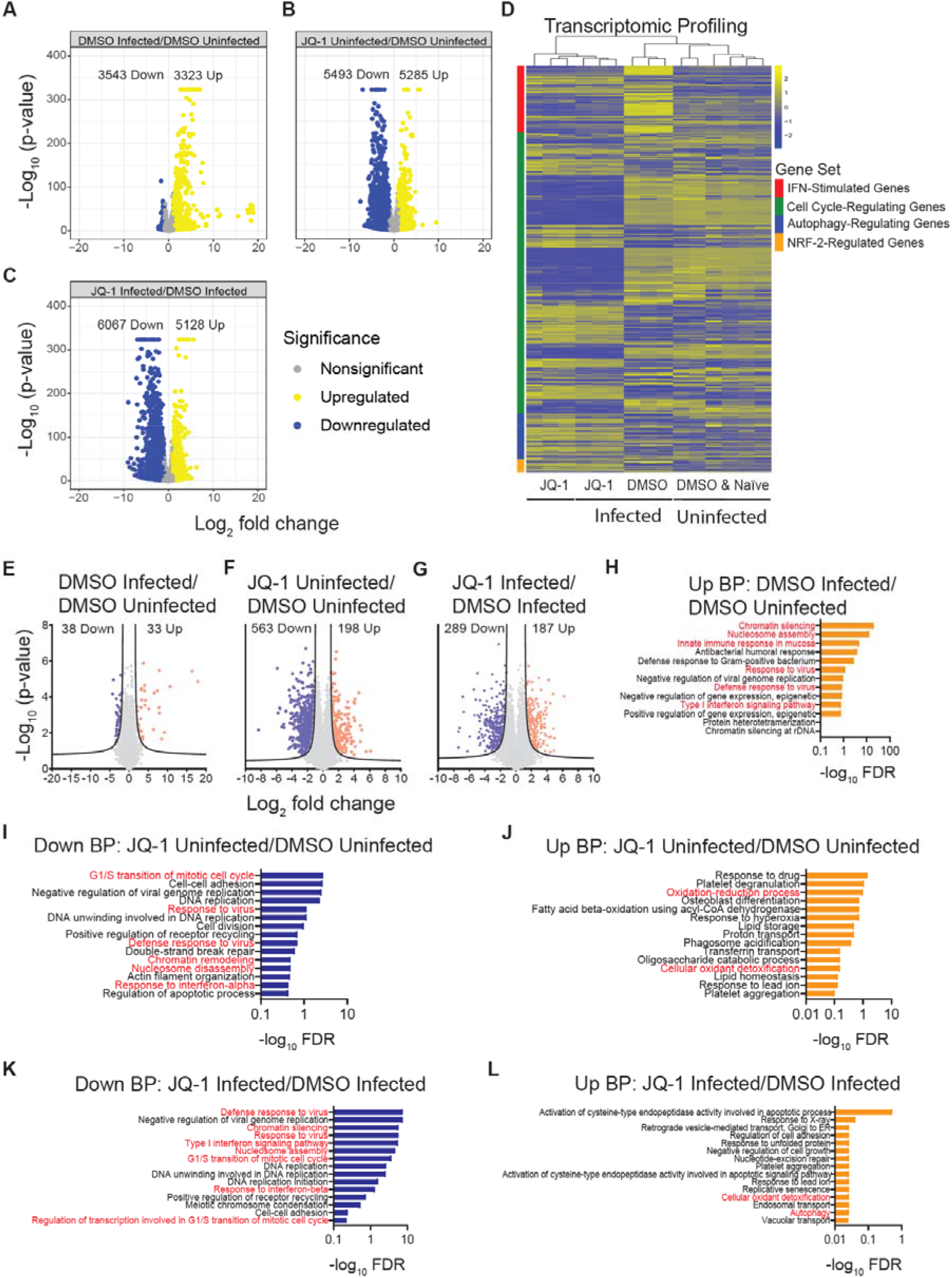
SARS-CoV-2 infection- and JQ-1-mediated modulations of the chromatin accessibility landscape underlie the transcriptomic and proteomic profiles in lung epithelial Calu-3 cells. Volcano plots showing relative log_2_FC and statistical significance of differentially expressed genes in (**A**) DMSO-treated infected versus DMSO-treated uninfected, (**B**) JQ-1-treated infected versus DMSO-treated infected and (**C**) JQ-1-treated uninfected versus DMSO-treated uninfected contrasts. The number of significant (FDR of ≤0.05) DRGs are indicated. (**D**) Heatmap of DRGs across four biological pathways (IFN-stimulated genes, cell cycle-regulating genes, autophagy-regulating genes, and NRF-2-regulated genes) scaled as z-score across rows. Volcano plots showing relative log_2_FC and statistical significance of differentially abundant proteins in (**E**) DMSO-treated infected versus DMSO-treated uninfected, (**F**) JQ-1-treated uninfected versus DMSO-treated uninfected and (**G**) JQ-1-treated infected versus DMSO-treated, infected contrasts. The number of significantly (FDR ≤0.05) regulated proteins is indicated. Top enriched (FDR ≤0.05) GO Biological Process terms of upregulated proteins in (**H**) DMSO-treated infected versus DMSO-treated, uninfected, and differentially regulated proteins in (**I-J**) JQ-1-treated, infected versus DMSO-treated infected and (**K**-**L**) JQ-1-treated uninfected versus DMSO-treated uninfected contrasts.

Hierarchical clustering of genes driving selected biological pathways revealed that SARS-CoV-2 infection induced upregulation of IFN-stimulated genes (Fig. 5D). JQ-1-treated groups, irrespective of their infection status, shared similar transcriptomic signatures for the displayed pathways, as did the uninfected DMSO-treated and naïve groups. Compared to other groups, JQ-1 treatment altered the cell cycle transcriptomic profile and upregulated genes driving autophagy and NRF-2-mediated cellular cytoprotective response, irrespective of the infection status (Fig. 5D). As expected, IFN and cytokine-mediated signalling pathways were upregulated in SARS-CoV-2-infected samples, while pathways driving oxidative phosphorylation were among the top downregulated pathways (Sup. Fig. 3B). Conversely, JQ-1 treatment in the presence of infection upregulated the pathways driving oxidative phosphorylation and downregulated the IFN and cytokine signalling pathways (Sup. Fig. 3B). These JQ-1-driven changes were less pronounced or even absent in the absence of infection (Sup. Fig. 3B). Nucleosome assembly, organisation, chromatin assembly, and other pathways associated with regulation of gene transcription were equally enriched across all analysed contrasts (Sup. Fig. 3B).

To generate proteomic profiles, we resolved proteins from the whole cell lysate on linear SDS-PAGE gels, visualised the proteins with Bio-Safe Coomassie Blue stain (Sup. Fig. 3C) and subjected them to liquid chromatography coupled with tandem mass spectrometry (LC-MS/MS) as described elsewhere (Mhlekude et al., 2021). PCA from the proteomic data demonstrated sample clustering according to their experimental groups, reflecting similar proteomic profiles between individual samples in each group (Sup. Fig. 3D). SARS-CoV-2 infection induced differential abundance of 71 proteins, with 33 upregulated and 38 downregulated (Fig. 5E). In the absence of infection, JQ-1 treatment induced differential regulation of 761 proteins, 189 of which were upregulated and 563 downregulated (Fig. 5F). JQ-1 treatment in the presence of SARS-CoV-2 infection induced differential regulation of 476 proteins, with 187 upregulated and 289 downregulated (Fig. 5G).

GO term analysis of the biological processes revealed that IFN and innate immune signalling driven by the STATs, IRFs, and NF-κB-1 transcription factors were among the upregulated pathways in SARS-CoV-2-infected samples, as compared to uninfected samples (Fig. 5H). Furthermore, chromatin silencing and nucleosome assembly were the top two highly-enriched pathways. Irrespective of the infection status, IFN signalling, chromatin silencing, nucleosome assembly, and G1/S transition of cell cycle were among the downregulated pathways in JQ-1-treated groups (Fig. 5 I-K). Paradoxically, nucleosome assembly and chromatin organisation-related pathways were upregulated at the transcriptomic levels between the JQ-1-treated groups irrespective of the infection status (Sup. Fig. 5B), but downregulated at the proteomic level (Fig. 5I-K). On the other hand, irrespective of the infection status, the NRF-2-driven oxidant detoxification pathway that forms part of the cellular antiviral cytoprotective response was among the upregulated pathways in the JQ-1-treated groups (Fig. 5J-L). Moreover, the autophagy pathway, which is reduced by SARS-CoV-2 (Gassen et al., 2021), was upregulated in the JQ-1-treated group in the presence of infection (Fig. 5L). Together, the transcriptomic and proteomic changes largely recapitulate the modulated biological pathways, which are driven by transcription factors whose binding motifs are enriched as determined by ATAC-seq.

### JQ-1 treatment antagonises SARS-CoV-2-mediated suppression of the antiviral NRF-2-mediated cytoprotective response

Despite the growing literature on the anti-SARS-CoV-2 activity of JQ-1 and other iBETs (Gilham et al., 2021; Mills et al., 2021; Samelson et al., 2022; Vann et al., 2022), TFs orchestrating transcriptional responses in the context of JQ-1-mediated anti-SARS-CoV-2 activity remain to be investigated. For this purpose, we performed global TF activity profiling using the bulk RNA-seq dataset, which was collected in parallel with ATAC-seq data to identify TFs with significantly regulated gene module scores across our experimental groups. SARS-CoV-2 infection of Calu-3 cells induced activity of the inflammatory TF families (STATs, IRFs, and NFκB-1) (Fig. 6A), whose signalling is associated with severe COVID-19 cases (Lam et al., 2021). JQ-1 treatment in the presence of SARS-CoV-2 infection suppressed the activity of these inflammatory TF families, suggesting an anti-inflammatory effect of JQ-1 (Fig. 6A). Interestingly, while SARS-CoV-2 infection induced NFκB-1-driven canonical signalling pathway, it strongly suppressed NFκB-2-driven noncanonical signalling pathway, reflecting a more precise SARS-CoV-2-mediated regulation of NFκB signalling pathways.

**Figure 6.**
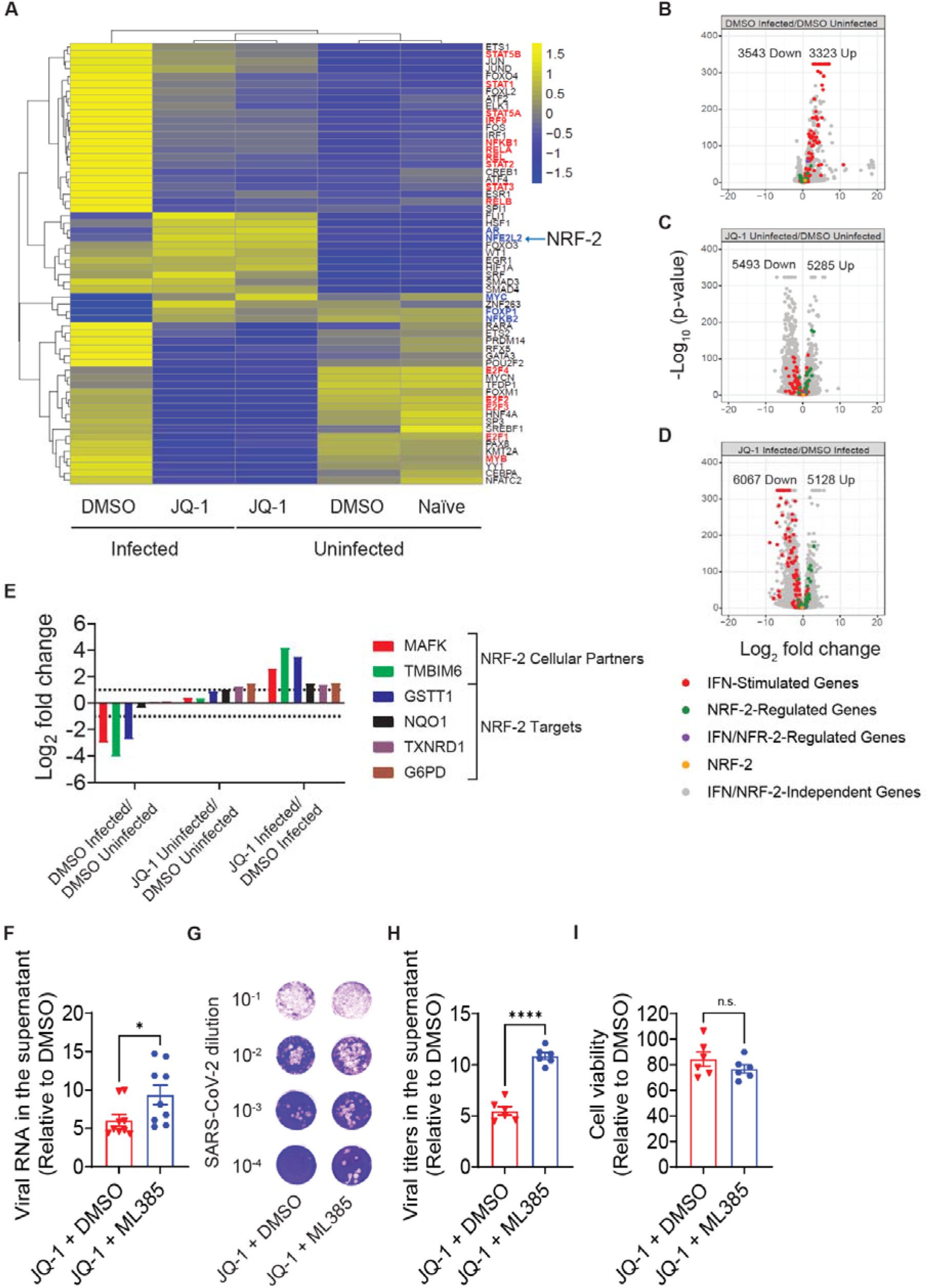
JQ-1 treatment antagonises SARS-CoV-2-mediated suppression of the antiviral NRF-2-mediated cytoprotective response. (**A**) Scaled TF activity scores in different treatments for RNAseq data, based on the Dorothea database. Volcano plots showing relative log_2_FC and statistical significance of differentially expressed genes involved in IFN and NRF-2 signalling in (**B**) DMSO-treated infected versus DMSO-treated uninfected, (**C**) JQ-1-treated infected versus DMSO-treated infected and (**D**) JQ-1-treated uninfected versus DMSO-treated uninfected contrasts. Log_2_FC analysis of differentially expressed proteins implicated in NRF-2 signalling. The bars indicate relative log_2_FC in expression between indicated contrasts and proteins with a relative log_2_FC of >1 and a FDR≤0.05 were considered significant. Relative quantification of (**F**) viral RNA copies and (**G-H**) infectivity from SARS-CoV-2 infected Calu-3 cells treated as indicated. The data were expressed as the normalised percentage infection relative to DMSO only-treated cells. Error bars show the SEM from triplicates of three and two independent experiments respectively. Cell viability analysis in Calu-3 cells treated as indicated. The graphs show background-subtracted and normalised data relative to DMSO only-treated cells. Error bars show the SEM from triplicates of two independent experiments. Unpaired nonparametric Student *t*-test was used to compare the means between experimental groups.

Irrespective of the infection status, JQ-1 treatment suppressed activity of the TF families that regulate cell cycle and proliferation (E2F & MYB), whose synergistic signalling is associated with severe lung injury in COVID-19 cases (Lam et al., 2021). This is in line with JQ-1-mediated dysregulation of cell cycle-regulating genes (Fig. 5D) and downregulation of pathways driving G1/S transition of the mitotic cell cycle captured in the proteomic data in (Figs. 5 I-K). Furthermore, JQ-1 also induced the transcriptional activities of nuclear factor (erythroid-derived-2)-like-2 (NFL2L2)/(NRF-2) and androgen receptor (AR), which drive a cytoprotective response (He et al., 2020) and RNA alternative splicing (Rana et al., 2021; Shah et al., 2020), respectively (Fig. 6A). It is noteworthy that pharmacological induction of NRF-2 signalling by 4-OI and DMF inhibits SARS-CoV-2 replication *in vitro* (Olagnier et al., 2020). NRF-2 and AR belong to CNC-bZIP and steroid nuclear receptor (NR) families of TFs, respectively; whose binding motifs were significantly enriched in the accessible ATAC-seq peaks from JQ-1-treated cells independent of the infection status (Sup. Table 1B-C). Together, these data suggest that the chromatin regulatory regions captured in our ATAC-seq data were not only accessible, but also transcriptionally active.

To discern between SARS-CoV-2-mediated upregulation of IFN signalling, inhibition of NRF-2 signalling, and JQ-1-mediated antagonistic effects in these processes, we generated volcano plots to analyse the log_2_FCs of STAT/IRF and NRF-2 target genes. As expected, IFN-stimulated genes (ISGs) were upregulated in the context of SARS-CoV-2 infection (Fig. 6B). SARS-CoV-2 infection showed minimal changes on the expression of NRF-2 target genes and no effect on NRF-2 expression *per se* (Fig. 6B). JQ-1 treatment induced the downregulation of ISGs in the absence (Fig. 6C) and presence (Fig. 6D) of SARS-CoV-2 infection. Irrespective of the infection status, JQ-1 treatment induced an upregulation of the expression of NRF-2 target genes (Fig. 6C-D). Like SARS-CoV-2 infection (Fig. 6B), JQ-1 treatment did not significantly change expression of NRF-2 itself, independent of the infection status (Fig. 6C-D). These data suggest that both SARS-CoV-2 and JQ-1-mediated modulations of NRF-2 signalling do not significantly alter NRF-2 expression *per se*, but likely modulate the expression of its signalling cofactors.

NRF-2 induces gene expression by binding to the *cis*-acting antioxidant response elements (ARE) on the promoters of its target genes as a heterodimer with the small musculoaponeurotic fibrosarcoma (MAF) transcription factors, which act as its indispensable cofactors (Itoh et al., 1997; Katsuoka and Yamamoto, 2016). Furthermore, transmembrane B cell lymphoma 2-associated X protein (BAX) inhibitor motif-containing 6 (TMBIM6) protects the host against oxidative stress by inducing NRF-2 signalling (Kim et al., 2020; Lee et al., 2007). Interestingly, the proteomic data showed significant reduction of log_2_FCs for MAFK and TMIBM6, and NRF-2 target proteins in SARS-CoV-2-infected Calu-3 cells (Fig. 6E). JQ-1 treatment antagonised these SARS-CoV-2-mediated changes, with more pronounced inhibition in the context of infection (Fig. 6E). These data suggest that SARS-CoV-2 suppresses NRF-2 signalling by downregulating its indispensable signalling cofactors.

Further log_2_FC analysis revealed that SARS-CoV-2 infection induced a dramatic upregulation of COQ6 expression (Sup. Fig. 4A), which drives the production of coenzyme Q10 to shuttle electrons between complexes I, II and II; and associated dehydrogenase enzymes in the mitochondrial electron transport chain (Acosta Lopez et al., 2019; Wang and Hekimi, 2016). These data hint at the need for a ubiquinone biosynthetic pathway in SARS-CoV-2 pathogenesis, despite the defective oxidative phosphorylation (Sup. Fig. 3B). Moreover, SARS-CoV-2 infection downregulated COX16 that serves as an indispensable component of cytochrome-c-oxidase (complex IV) in the electron transport chain (Wintjes et al., 2021) and upregulated the chromatin silencing histone variants (H2AFY, H2AFV and H2AFZ) (Sup. Fig. 4A), which likely drive the SARS-CoV-2-mediated upregulation of chromatin silencing and nucleosome assembly pathways (Fig. 5H). More importantly, SARS-CoV-2 downregulated the expression of YTHDF2, a methyl-6-adenosine (m6A) reader of the RNA molecules in the host that binds to and degrade m6A-bearing host (Du et al., 2016) and viral (Gokhale et al., 2016; Gonzales-van Horn and Sarnow, 2017; Lichinchi et al., 2016) RNA molecules to prevent the expression of aberrant cellular RNA and inhibit viral replication, respectively. JQ-1 displayed an antagonistic effect against the above-mentioned SARS-CoV-2-mediated proteomic modulations (Sup. Fig. 4A).

NRF-2 signalling, which is induced by JQ-1 treatment in Calu-3 cells (Fig. 6A), inhibits SARS-CoV-2 infection when induced by 4-OI and DMF (Olagnier et al., 2020). This prompted us to investigate whether pharmacological inhibition of NRF-2 signalling antagonises JQ-1-mediated anti-SARS-CoV-2 activity. Interestingly, co-treating Calu-3 cells with JQ-1 and ML385, a specific inhibitor of NRF-2 (Singh et al., 2016), prior to infection, led to a significant increase in viral RNA copies (Fig. 6F) and infectious titers (Fig. 6G-H) in the supernatant compared to JQ-1 and DMSO co-treated cells, in the absence of detectable cell toxicity (Fig. 7I). These data suggest that JQ-1 inhibition of SARS-CoV-2 infection in Calu-3 cells involves efficient NRF-2 signalling. Together, these data shed light on JQ-1-mediated multi-pronged approach underlying its anti-SARS-CoV-2 activity.

**Figure 7.**
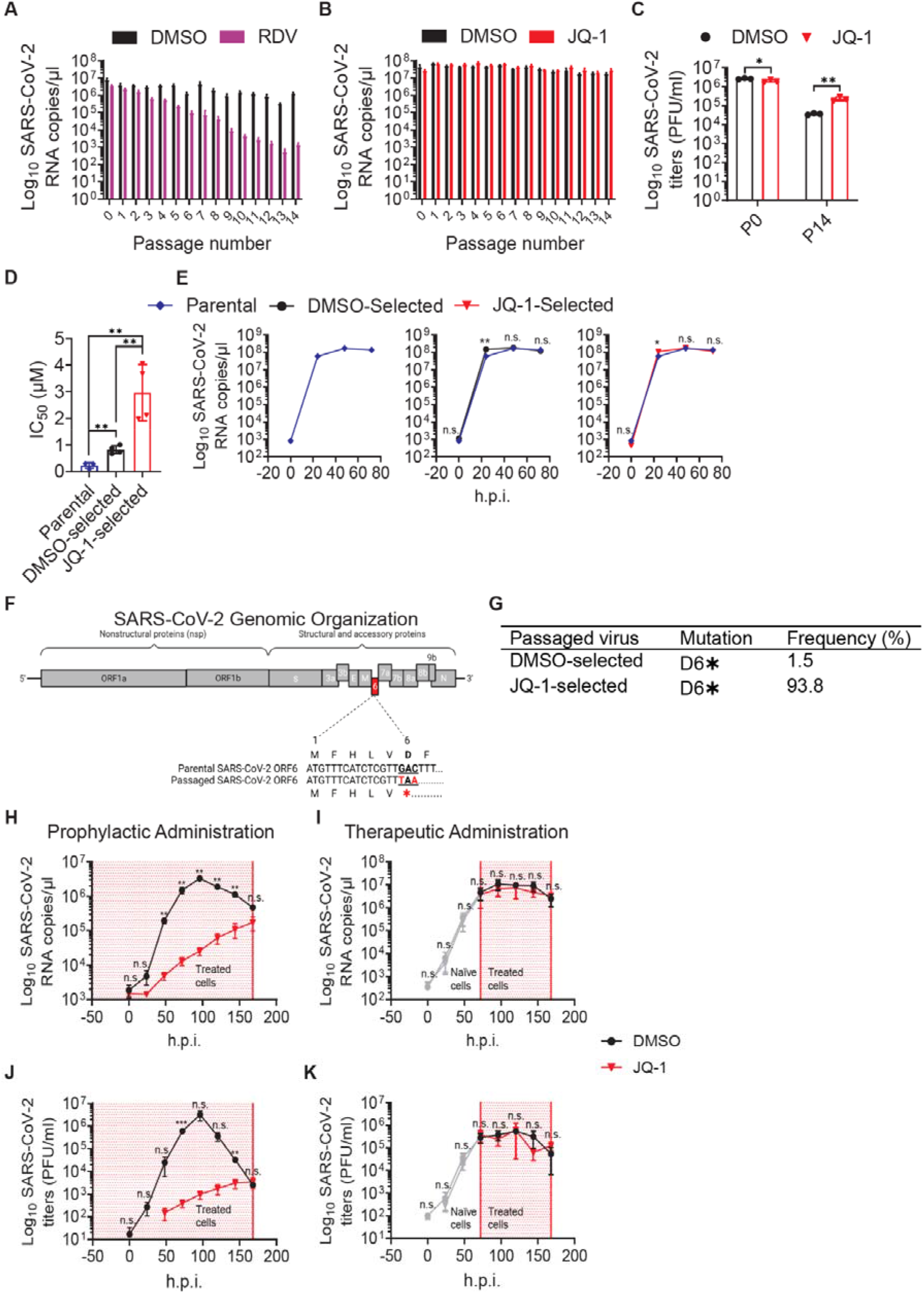
SARS-CoV-2 infection subverts JQ-1-mediated antiviral state in lung epithelial Calu-3 and human bronchial airway epithelial cells (hBAECs). Quantification of SARS-CoV-2 RNA (copies/μl) from triplicates of 15 serial passages under two-fold escalating concentrations of (**A**) remdesivir and (**B**) JQ-1, with DMSO as a mock. **(C)** Quantification of SARS-CoV-2 titers (PFU/ml) in triplicates from P1 (JQ-1 0.32μM) and P14 (JQ-1 5.12μM) passages. The means between experimental groups were compared using nonparametric Student *t*-test. Quantification of IC_50_ values from JQ-1 dose-dependent inhibition curves determined in JQ-1-treated Calu-3 cells infected with parental and passaged SARS-CoV-2 virions (MOI 0.1). The error bars represent the mean±SEM from four independent experiments and the means between experimental groups were compared using nonparametric Student *t*-test. Viral growth kinetics in DMSO-treated Calu-3 cells showing viral RNA quantities from parental and passaged (P14) SARS-CoV-2 virions (MOI 0.1) from three independent experiments. The means between experimental groups were compared using nonparametric multiple *t*-test. Schematic diagram of the SARS-CoV-2 genome depicting the mutation creating a premature stop codon at position six of ORF6 among the passaged (P14) virions. **(G)** Quantification of the ORF6 D6✱ mutation frequency from sequences from passaged (P14) SARS-CoV-2 virions. Virus growth kinetics in DMSO and JQ-1 (2.56 μM)-treated hBAECs depicting SARS-CoV-2 RNA quantities (copies/μl) in the contexts of (**H**) prophylactic and (**I**) therapeutic drug administration from three independent experiments. Virus growth kinetics in DMSO and JQ-1 (2.56 μM)-treated hBAECs depicting SARS-CoV-2 titers (PFU/ml) in the contexts of (**J**) prophylactic and (**K**) therapeutic drug administration from three independent experiments.

### SARS-CoV-2 subverts JQ-1-mediated antiviral state in lung epithelial Calu-3 and human bronchial airway epithelial cells

SARS-CoV-2 can adapt to and acquire resistance against potent virus-directed antivirals such as remdesivir (RDV) *in vitro* (Stevens et al., 2022; Szemiel et al., 2021), a replication bottleneck that induces the emergence SARS-CoV-2 variants *in vivo* (Heyer et al., 2022). To investigate whether SARS-CoV-2 can acquire resistance to iBETs, we serially passaged it in Calu-3 cells under escalating two-fold concentrations of JQ-1 (Sup. Fig. 5A), along with RDV as a reference. SARS-CoV-2 serial passaging under increasing two-fold concentrations of RDV led to a gradual reduction of viral RNA copies in the supernatant, with a potential resistance phenotype emerging at the earliest at passage 14 (Fig. 7A). In contrast, SARS-CoV-2 serial passaging under escalating two-fold concentrations of JQ-1 displayed unrelenting high concentrations of viral RNA copies from passage two until passage 14. These data suggest a transient nature of JQ-1-mediated anti-SARS-CoV-2 activity and early adaptation of SARS-CoV-2 to JQ-1 (Fig. 7B). Quantification of viral titers from passages one and 14 revealed that SARS-CoV-2 virions were sensitive to JQ-1 treatment at the beginning of the passaging experiments, and acquired resistance and replication fitness to JQ-1 during serial passaging (Fig. 7C).

To determine the extent to which SARS-CoV-2 passaging under JQ-1 led to the acquisition of resistance, we investigated the sensitivity of DMSO- and JQ-1-selected SARS-CoV-2 virions to JQ-1 in Calu-3 cells by quantifying the viral RNA copies in the supernatant and calculating IC_50_ concentrations. Compared to parental SARS-CoV-2, virions that were serially passaged under JQ-1 exhibited reduced sensitivity to JQ-1 treatment in Calu-3 cells (Fig. 7D), which was more pronounced (12-fold) compared to DMSO-selected virions (4-fold) (Sup. Fig. 5B). Viral growth kinetics in Calu-3 cells showed that DMSO- and JQ-1-selected SARS-CoV-2 virions may have acquired some degree of replication fitness compared to the parental SARS-CoV-2 virions (Fig. 7E). Next-Generation Sequencing (NGS) revealed virus sequences encoding a premature stop codon at position six of ORF6 in the genome from virions passagaged in the presence of JQ-1, as opposed to parental virus (Fig. 7F). This premature stop codon was present far more frequently in JQ-1-selected virions (93.8% of reads covering this region) than DMSO-selected virions (1.5% of reads covering this region) (Fig. 7G).

Finally, we performed kinetic, multi-cycle infection experiments in hBAECs to compare JQ-1-mediated anti-SARS-CoV-2 activity in the contexts of prophylactic and therapeutic treatment. In the set-up mimicking prophylactic administration, we started treatment of hBAECs with DMSO or JQ-1 48 hours prior to SARS-CoV-2 infection (Sup. Fig. 5C). For therapeutic administration, we infected naïve hBAECs and started treatment 48 hours post-infection (Sup. Fig. 5D). JQ-1 treatment was continued for the duration of the entire experiment. Prophylactic administration of JQ-1 resulted in antiviral activity that was more pronounced earlier post infection, which however vanished over time, as indicated by the gradual increase in viral RNA copies (Fig. 7H) and infectious titers (Fig. 7J). Strikingly, administration of JQ-1 post establishment of infection failed to curb SARS-CoV-2 replication, as indicated by similar viral RNA copies (Fig. 7I) and infectious titers (Fig. 7I) between DMSO- and JQ-1-treated groups. Together, these data suggest that JQ-1 treatment results in a transient antiviral state, which however can be subverted by SARS-CoV-2.

The area shaded in red indicates the time period of drug administration and the grey lines in the context of therapeutic administration indicate infection before drug administration. The means between experimental groups were compared using nonparametric multiple *t*-test. Error bars represent the mean±SEM.

## DISCUSSION

Prior studies attributed the iBET-mediated anti-SARS-CoV-2 activity to the downregulation of cellular ACE2 expression (Gilham et al., 2021; Mills et al., 2021; Samelson et al., 2022), but failed to capture the pleiotropic nature of the antiviral state induced by these compounds and their susceptibility to SARS-CoV-2 subversion. Here, we show that JQ-1-mediated anti-SARS-CoV-2 activity goes beyond the downregulation of ACE2 expression by inducing a vast transcriptional remodulation, including the mounting of antiviral NRF-2-mediated responses. Furthermore, we show that JQ-1-mediated antiviral activity is transient when administered prophylactically and nullified by SARS-CoV-2 when administered therapeutically following an established infection, suggesting effective viral antagonistic strategies.

In agreement with prior studies in Calu-3 cells (Gilham et al., 2021; Mills et al., 2021; Samelson et al., 2022; Vann et al., 2022), iBETs displayed an antiviral activity against SARS-CoV-2 infection in lung epithelial Calu-3 cells and hBAECs when administered prophylactically, suggesting that the iBET-mediated anti-SARS-CoV-2 activity is readily translated from cell lines *in vitro* to clinically relevant human-derived infection models. The antiviral activity of JQ-1 and its analogues has largely been reported in the context of SARS-CoV-2 parental strains (Gilham et al., 2021; Mills et al., 2021; Samelson et al., 2022) and VOC Delta (Vann et al., 2022). Our data showing sensitivity of SARS-CoV-2 and SARS-CoV, but not MERS-CoV, to JQ-1 suggest that MERS-CoV replication, and likely that of other *Merbecoviruses*, is independent of BET proteins. This difference can potentially be attributed to the interactions between SARS-CoV-2-E and cellular BET proteins (Chen et al., 2022; Vann et al., 2022), through a conserved histone motif present in SARS-CoV-E and SARS-CoV-2-E proteins (Gordon et al., 2020; Schoeman and Fielding, 2020), which is lacking in MERS-CoV-E protein (Schoeman and Fielding, 2020).

The regulatory landscape orchestrating the iBET-mediated transcriptional responses to SARS-CoV-2 infection has remained largely unexplored. Accessible peaks in SARS-CoV-2-infected samples were enriched with binding motifs for the inflammatory TF family (STATs, IRFs and NFκB), which were significantly associated with the upregulation of IFN and cytokine signalling pathways. On the other hand, accessible peaks from the same samples were significantly associated with the downregulation of smell receptor signalling pathways, reminiscent of the olfactory dysfunction reported in COVID-19 cases (Brann et al., 2020; de Melo et al., 2021; Qiu et al., 2020). Significant association of accessible peaks from JQ-1-treated samples in the presence of infection with the downregulation of the smell receptor signalling pathways suggests that JQ-1 is unable to dampen SARS-CoV-2-mediated olfactory dysfunction (Brann et al., 2020; de Melo et al., 2021; Qiu et al., 2020).

Similar to the ATAC-seq data, we captured massive differential regulation of genes and proteins in JQ-1-treated samples, irrespective of infection, compared to SARS-CoV-2 infection. Upregulated genes and proteins in SARS-CoV-2-infected samples were significantly associated with the upregulation of IFN and cytokine signalling pathways. In line with the detected enrichment of accessible binding motifs for the inflammatory TF family, supervised hierarchical clustering further showed that SARS-CoV-2 infection induced the transcription of IFN-stimulated genes. In accordance with reported SARS-CoV-2-mediated impairment of mitochondrial biogenesis (Guarnieri et al., 2022), downregulated genes in SARS-CoV-2-infected samples were significantly associated (among others) with the downregulation of pathways driving mitochondrial oxidative phosphorylation (OXPHOS). The upregulation of chromatin silencing and nucleosome assembly pathways at the proteomic level in SARS-CoV-2-infected samples is consistent with SARS-CoV-2-ORF8-mediated chromatin compaction (Kee et al., 2021) and may represent the epigenomic mechanism underlying SARS-CoV-2-mediated host-shutoff (Finkel et al., 2021).

Interestingly, despite inducing a strong NFκB-1 TF activity that drives inflammatory cytokine production (Ramasamy and Subbian, 2021), SARS-CoV-2 infection induced an equally strong suppression of NFκB-2 TF activity, which regulates the maturation of antibody-producing B cells and development of germinal centres (GC) in the lymph nodes (Caamaño et al., 1998; Silva et al., 2016). Conversely, irrespective of infection, JQ-1 suppressed NFκB-1 TF activity and induced NFκB-2 TF activity. It is noteworthy that inhibition of NFκB-1 transcriptional footprint inhibits SARS-CoV-2 replication (Nilsson-Payant et al., 2021). These data reflect a carefully executed and precise regulation of NFκB-mediated signalling pathways by SARS-CoV-2 infection and JQ-1 administration.

Association of the upregulated genes and proteins in JQ-1-treated samples with the downregulation of IFN signalling and viral genome replication pathways validates JQ-1-mediated suppression of innate immune responses (Chen et al., 2022; Mills et al., 2021; Samelson et al., 2022) and antiviral activity (Gilham et al., 2021; Mills et al., 2021; Samelson et al., 2022; Vann et al., 2022). SARS-CoV-2 infection limits induction of autophagy (Gassen et al., 2021), and pharmacological induction of NRF-2, which is a cellular cytoprotective response-inducing transcription factor that belongs to CNC-bZIP TF family, inhibits SARS-CoV-2 replication (Olagnier et al., 2020). Of note, a binding motif for CNC-bZIP TF family was among the top enriched TF motifs in accessible peaks from JQ-1-treated samples, irrespective of infection.

Of particular interest, supervised hierarchical clustering revealed that JQ-1 treatment upregulated the expression of autophagy-regulating and NRF-2 target genes, irrespective of SARS-CoV-2 infection. In accordance, pathway enrichment analysis at the proteomic level revealed upregulation of autophagy in JQ-1-treated samples in the presence of infection and NRF-2-mediated cellular oxidant detoxification pathway irrespective of infection status. Motif enrichment for the TFs belonging to CNC-bZIP and NFY families in the context of JQ-1 administration, irrespective of infection, revealed the chromatin regulatory landscape underlying JQ-1-mediated induction of NRF-2 signalling (Lv et al., 2022), which exhibits an anti-SARS-CoV-2 activity in the context of DMF- and 4-OI-mediated induction (Olagnier et al., 2020). Moreover, signalling by the NFY TFs increases chromatin accessibility by preventing nucleosome encroachment (Oldfield et al., 2014, 2019). Therefore, enrichment of the binding motifs for the NFY TFs supports the highly accessible chromatin landscape detected in the context of JQ-1 treatment. Together, these data show that JQ-1 induced the activities of NRF-2, which drive the induction of antiviral cellular cytoprotective response (Olagnier et al., 2020).

SARS-CoV-2 infection upregulated IFN-stimulated genes, but displayed minimal alterations on the expression of NRF-2 target genes, without altering NRF-2 expression *per se*. On the other hand, JQ-1 administration downregulated the expression of IFN-stimulated genes and induced the expression of NRF-2 target genes irrespective of infection. However, like SARS-CoV-2 infection, JQ-1 administration also did not alter NRF-2 expression, despite significantly altering the expression of its target genes. These data suggest that differential regulation of NRF-2 signalling by SARS-CoV-2 infection and JQ-1 administration does not alter NRF-2 expression *per se*, but likely the expression of its signalling co-factors.

Accordingly, the proteomic data showed that SARS-CoV-2 infection induced significant log_2_FC downregulation of MAFK and TMBIM6, which act as indispensable co-factors in the induction of NRF-2 signalling (Itoh et al., 1997; Katsuoka and Yamamoto, 2016; Kim et al., 2020; Lee et al., 2007). Accompanying SARS-CoV-2-mediated downregulation of NRF-2 signalling co-factors was the downregulation of NRF-2 target proteins. JQ-1 administration induced an upregulation of MAFK, TMBIM6, and NRF-2 target proteins, with a more pronounced effect in the presence of infection. These data complement SARS-CoV-2- and JQ-1-mediated differential regulation of NRF-2 TF activity and suggest that both SARS-CoV-2 infection and JQ-1 administration regulate NRF-2 TF activity by modulating the expression of its signalling co-factors and not NRF-2 expression *per se*. Suppression of NRF-2-mediated cytoprotective response in biopsies from COVID-19 patients (Olagnier et al., 2020) supports the biological plausibility of our data.

DMF- and 4-OI-mediated induction of NRF-2-driven cytoprotective protective response with anti-SARS-CoV-2 activity (Olagnier et al., 2020) and known JQ-1-mediated induction of NRF-2 signalling (Lv et al., 2022), prompted us to pursue the effect of JQ-1-mediated induction of NRF-2 signalling in the context of JQ-1-mediated anti-SARS-CoV-2 activity. Partial antagonism of JQ-1-mediated anti-SARS-CoV-2 activity by ML385, a specific inhibitor of NRF-2 (Singh et al., 2016), suggest that induction of NRF-2 signalling by JQ-1 administration contributes to JQ-1-mediated anti-SARS-CoV-2 activity, which is consistent with NRF-2-mediated anti-SARS-CoV-2 reported in the context DMF- and 4-OI-mediated induction (Olagnier et al., 2020). The combination of JQ-1-mediated downregulation of ACE2 expression and induction of NRF-2 signalling shown in our study, suggest that JQ-1 exhibits a pleiotropic anti-SARS-CoV-2 activity that affects multiple steps of the viral replication cycle.

How exactly JQ-1-mediated induction of NRF-2 signalling exerts an anti-SARS-CoV-2 activity remains unknown. Olagnier et al. proposed that the induction of hypoxia-inducible-factor-1-alpha (HIF-1α) gene expression program induced by 4-OI administration, which was downregulated in biopsies from COVID-19 patients along with NRF-2 signalling pathway, potentially contribute to NRF-2-mediated antiviral cytoprotective response (Olagnier et al., 2020). In contrast, Codo. et al. reported that SARS-CoV-2-mediated induction of ROS stabilises HIF-1α to induce its gene expression program that promotes glycolysis to sustain SARS-CoV-2 replication in monocytes (Codo et al., 2020). Consistent with the later report, we found induction of HIF-1α TF activity following SARS-CoV-2 infection in Calu-3 cells.

Furthermore, irrespective of infection, JQ-1 administration also induced HIF-1α TF activity. Downregulation of HIF-1α TF activity in control groups suggest that both SARS-CoV-2 infection and JQ-1 administration induce HIF-1α signalling. This is consistent with the idea of alternative JQ-1-driven HIF-1α TF activity-inducing mechanisms that are independent of SARS-CoV-2-mediated ROS production and argues against the potential effect of HIF-1α-mediated signalling in NRF-2-induced anti-SARS-CoV-2 activity proposed elsewhere (Olagnier et al., 2020). Alternatively, NRF-2 signalling induces the expression of a cytoprotective enzyme called heme-oxygenase-1 (HO-1) (Olagnier et al., 2020; Wagener et al., 2020), which catalyses the catabolism of heme into biliverdin, iron, and carbon monoxide (CO) (Dunn et al., 2014). Interestingly, the products of HO-1-mediated heme catabolism exhibit a broad-spectrum antiviral activity (Espinoza et al., 2017).

Of particular interest, prophylactic administration of free iron and biliverdin to Vero E6 cells prior to infection inhibited SARS-CoV-2 replication (Kim et al., 2021), In contrast, despite binding to SARS-CoV-2-S protein with nanomolar affinity and dampening spike interactions with neutralising antibodies, administration of 100μM biliverdin at the time of infection did not inhibit infection of Vero E6 cells by SARS-CoV-2 (Rosa et al., 2021). These studies (Kim et al., 2021; Rosa et al., 2021) suggest that like JQ-1, biliverdin-induced anti-SARS-CoV-2 activity is directed at the host and not the infecting virions. This is consistent with the maintenance of infection efficiency by the virions secreted under JQ-1 treatment. Taken together, these studies highlight the mechanism underlying NRF-2-mediated anti-SARS-CoV-2 activity, which itself remains to be fully unravelled.

Immune selection of SARS-CoV-2 has led to the emergence of VOCs bearing an arsenal of mutations in the spike and other viral proteins, which confer immune evasion capabilities and transmission superiority (Tuekprakhon et al., 2022; Willett et al., 2022). SARS-CoV-2 adaptation to virus-directed antivirals such as remdesivir (RDV) has been reported *in vitro* (Stevens et al., 2022; Szemiel et al., 2021) and *in vivo* (Heyer et al., 2022). Here, as early as passage one (P1) under JQ-1 selection, SARS-CoV-2 RNA copies and infectious titers increased in the supernatant and reached a plateau that was maintained across 14 passages despite the subsequent two-fold escalations of JQ-1 concentrations. Compared to the parental strain, JQ-1-selected, and to a lesser extent also DMSO-selected, SARS-CoV-2 virions showed reduced sensitivity to JQ-1 treatment in Calu-3 cells. These data suggest that passaging under DMSO, and even more under JQ-1, induced the emergence of SARS-CoV-2 variants with enhanced viral fitness and reduced sensitivity to JQ-1, unlike the acquisition of resistance to RDV that led to viral replication defect *in vitro* (Stevens et al., 2022; Szemiel et al., 2021).

NGS analysis of SARS-CoV-2 virions passaged under JQ-1 treatment revealed acquisition of a premature stop codon corresponding to position six of ORF6, (ORF6^D6STOP^ mutation), a mutation that was less pronounced in DMSO-selected virions. Like JQ-1, SARS-CoV-2-ORF6 antagonises IFN signalling in infected cells (Miorin et al., 2020; Schroeder et al., 2021; Xia et al., 2020). This IFN signalling-repressive effect of JQ-1 may have minimised the need for ORF6-mediated repression of IFN signalling and promoted ORF6 trade-offs. This is consistent with the early emergence of SARS-CoV-2 variants harbouring an in-frame deletion in ORF6 following passaging in IFN-deficient Vero E6 cells (Riojas et al., 2020). In contrast, replication of virions passaged under DMSO are expected to benefit from preserving the ORF6-mediated antagonism of IFN signalling. Investigation of how the acquisition of ORF6^D6STOP^ mutation mediates resistance to JQ-1-mediated antiviral activity and whether this ORF6 trade-off constitutes an antagonistic pleiotropy by enhancing sensitivity to IFN treatment is warranted. Together, these data suggest that acquisition of the ORF6^D6STOP^ mutation is not uniquely induced by JQ-1 treatment, but gains more dominance after encountering an IFN-deficient milieu induced by JQ-1 treatment.

The accumulating evidence from our study and previous studies (Gilham et al., 2021; Mills et al., 2021; Samelson et al., 2022; Vann et al., 2022) show that iBETs exhibit an anti-SARS-CoV-2 activity when administered prophylactically and not therapeutically (Chen et al., 2022). Whether SARS-CoV-2 infection counteracts JQ-1-mediated infection barriers or acquires alternative capabilities to infect and replicate in the host remains to be addressed in future studies. Interestingly, in one study (Chen et al., 2022), administration of iBETs at the time of infection, as opposed to pre-treatment, resulted in absence of an inhibitory effect and even exacerbation of SARS-CoV-2 replication. In line with this report, JQ-1 displayed no antiviral activity when administered in a therapeutic set-up in our study. The proviral effect of therapeutically administered JQ-1 reported by (Chen et al., 2022), which is absent in our study, may be related to different time points of sample harvest following infection (48 versus 24 hours, respectively). Since SARS-CoV-2 infection potently suppresses NRF-2-mediated antiviral cytoprotection responses (Olagnier et al., 2020), subsequent and long-term JQ-1 administration may be unable to revert this effect, respectively, resulting in SARS-CoV-2-mediated nullification of JQ-1-mediated anti-SARS-CoV-2 activity. Together, these data evoke questions about the clinical suitability for both prophylactic and therapeutic administration of JQ-1 (and likely other iBETs) in the context of COVID-19 and illuminate the potential hurdles that iBETs will have to overcome in order to improve disease prognosis.

## ACKNOWLEDGEMENTS

We thank J. S. M. Peiris, University of Hong Kong, China for providing the SARS-CoV isolate HKU-39849. We thank Victor Corman and his team for assistance with sequencing of virus stocks. We thank Dr. Tatiana A. Borodina for access to the MDC/BIH Genomic Core Facility. Computation has been performed on the HPC for Research/Clinic cluster of the Berlin Institute of Health. Parts of this work were supported by grants from NIAID-NIH CEIRS contract HHSN272201400008C to TCJ. This work was supported by funding of Berlin Institute of Health (BIH), Berliner SparkassenStiftung, Charité Stiftung and the DFG (SPP1923) to C.G., by Organo-Strat to C.G. and C.D. and German Center for Lung Research (DZL;82DZL002A1) to U.M. and R.O.

## AUTHOR CONTRIBUTIONS

B. Mhle. and C.G. conceived the study and designed the experiments. B. Mhle., S.Sten., J.J., A.R., J.H. and N.H. performed the experiments. R.O., S. Schr., U.M. and D.N. provided the material. B. Mhle., D.P., J.M.W., F.Z., B. Mue., T.C.J. and C.G. analysed the data. B. Mhle. and C.G. drafted the manuscript. M.A.M., C.D., A.P., V.T., D.N., G.S., D.B., and C.G. supervised the research. All authors reviewed and edited the manuscript.

