## Supplementary Figures for "Pharmacological inhibition of bromodomain and extra-terminal proteins induces NRF-2-mediated inhibition of SARS-CoV-2 replication and is subject to viral antagonism"

**Supplementary Material:**

**Supplementary Figures 1-5**

**Supplementary Table 1**

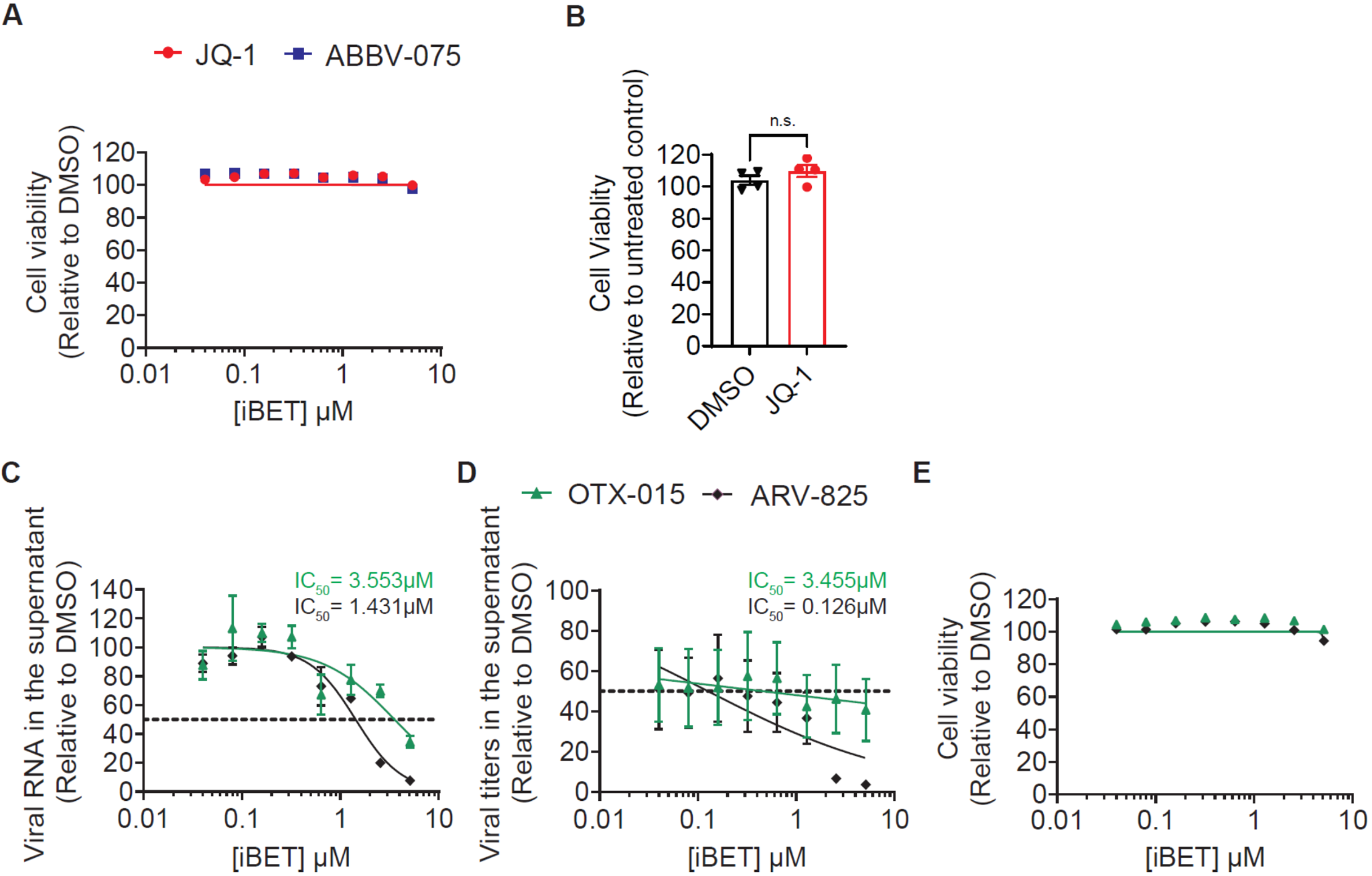

**Supplementary Figure 1: iBETs inhibit SARS-CoV-2 infection in lung epithelial Calu-3 cells.**

(**A**) Calu-3 cells were treated for 48 hours with indicated concentrations of iBETs and analysed for viability in three independent experiments.

(**B**) hBAECs treated with 2.56 µM JQ-1 for 48 hours were analysed for viability using duplicates of two independent experiments. The graphs show normalised data relative to DMSO treatment.

(**C**-**E**) Analysis of the antiviral potency of OTX-015 and ARV-825 in Calu-3 cells. Cells were pretreated with indicated concentrations of iBETs for 48 hours prior to infection with SARS-CoV-2 (MOI 0.1). Shown are DMSO-normalised (**C**) viral RNA abundance and (**D**) infectivity in supernatants from three independent experiments. (**E**) Relative viability was quantified luminometrically. Unpaired nonparametric Student *t*-test was used to compare the means between the experimental groups. Error bars represent the arithmetic mean±SEM.

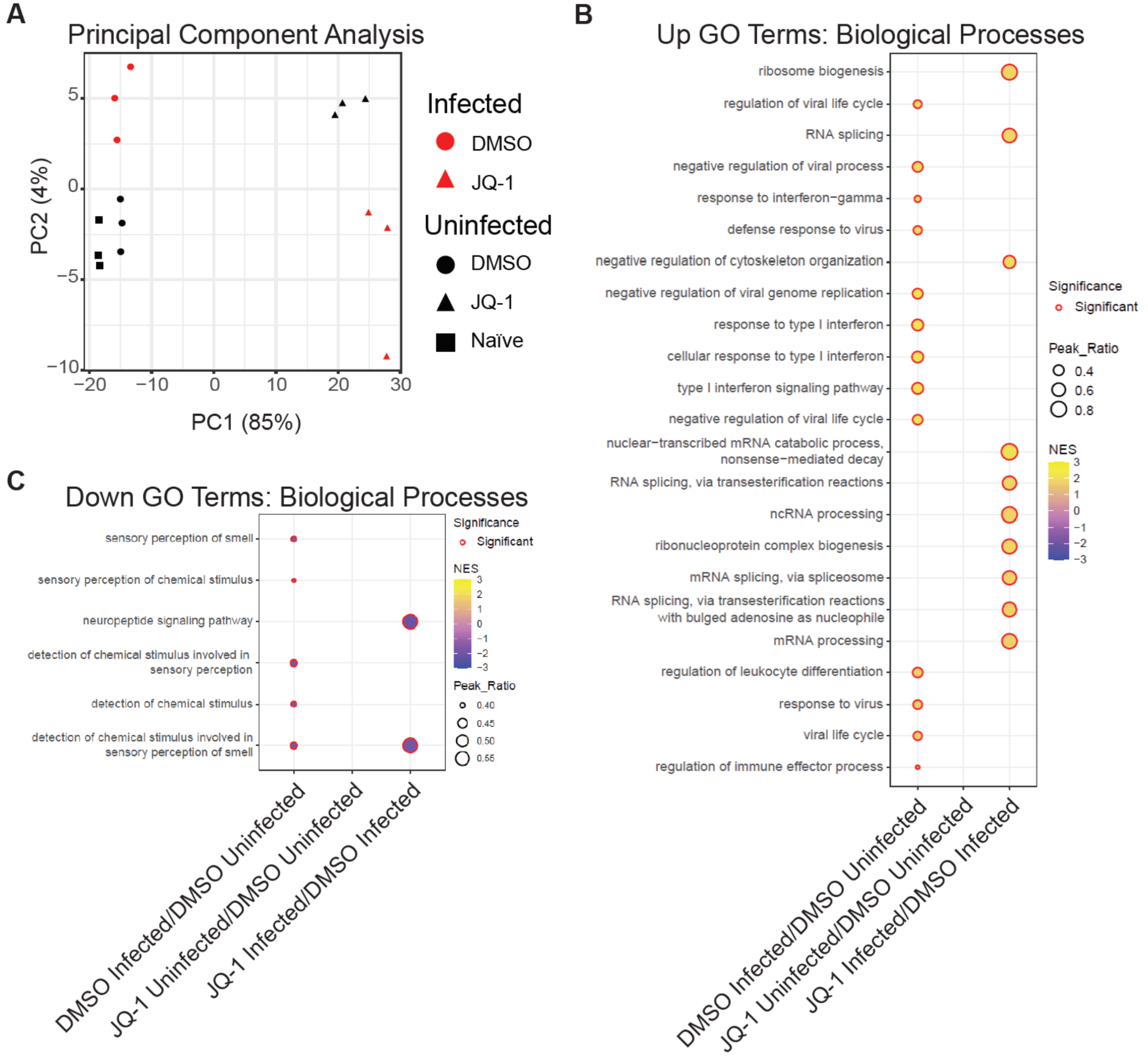

**Supplementary Figure 2. SARS-CoV-2 infection- and JQ-1-mediated modulations of the chromatin accessibility landscape modulate biological pathways in epithelial lung Calu-3 cells.**

(**A**) Principal Component Analysis of differentially accessible ATAC-seq peaks showing clustering of individual samples between experimental groups.

Dot plot showing gene ontology (GO Biological Process) analysis of genes annotated to (**B**) upregulated and (**C**) downregulated accessible ATAC-seq peaks between indicated contrasts. Dot colour represents normalised enrichment score (NES) and dot size (peak ratio) represents the number of significant peaks related to the GO term relative to the total number of significant peaks.

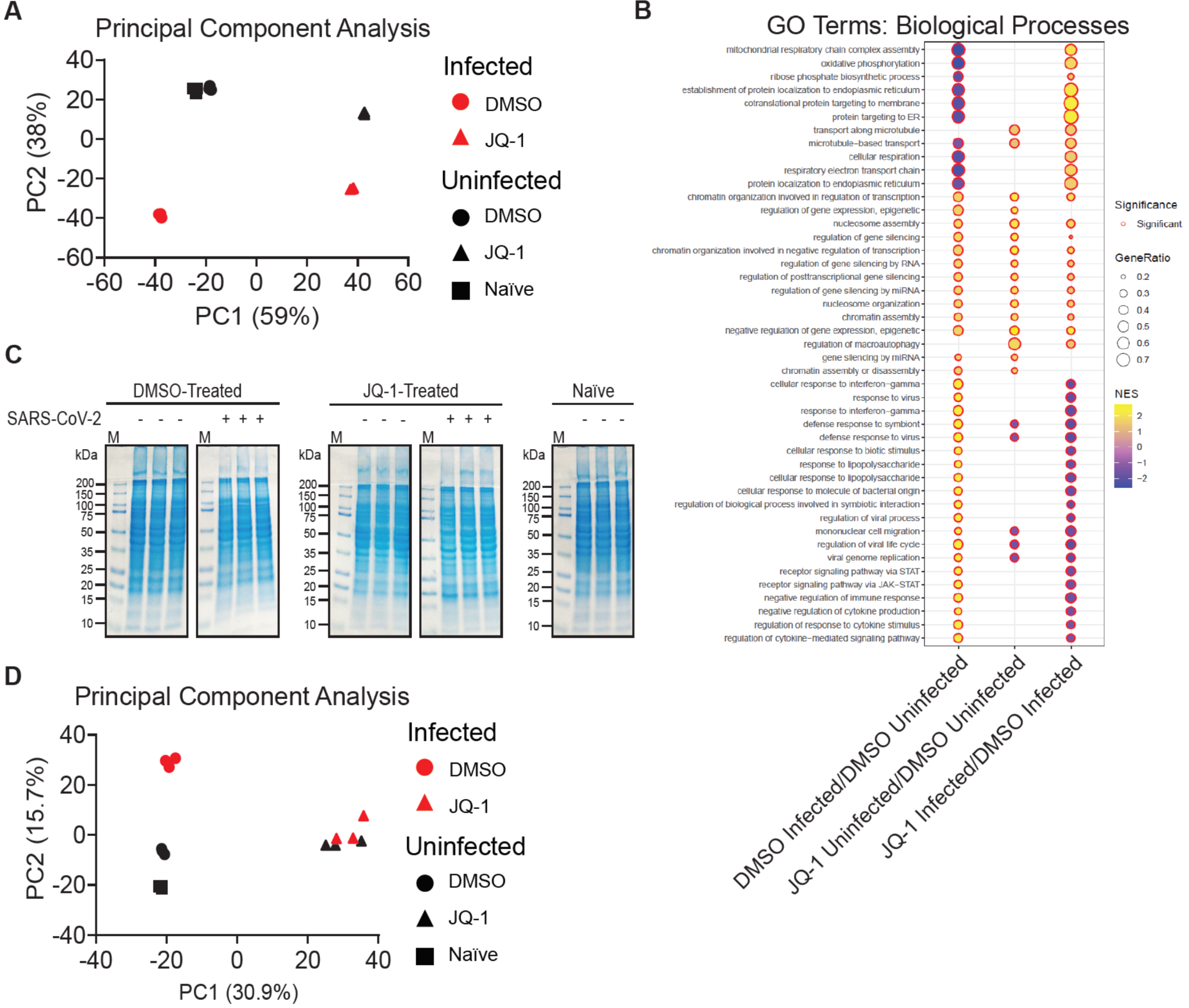

**Supplementary Figure 3. SARS-CoV-2 infection and JQ-1 treatment modulate the transcriptomic and proteomic profiles in epithelial lung Calu-3 cells**

(**A**) PCAs of DRGs showing clustering of individual samples between experimental groups.

(**B**) Dot plot showing gene ontology (GO Biological Process) analysis of DRGs between the selected contrasts. The dot colour and size represent normalised enrichment score (NES) and the number of genes related to the GO term relative to the total number of significant genes, respectively.

(**C**) Bio-Safe Coomassie Blue staining of proteins from Calu-3 cell lysates resolved on linear (7.5%) SDS-PAGE gels prior to mass spectrometry analysis.

(**D**) PCA of differentially abundant proteins showing clustering of individual samples between experimental groups.

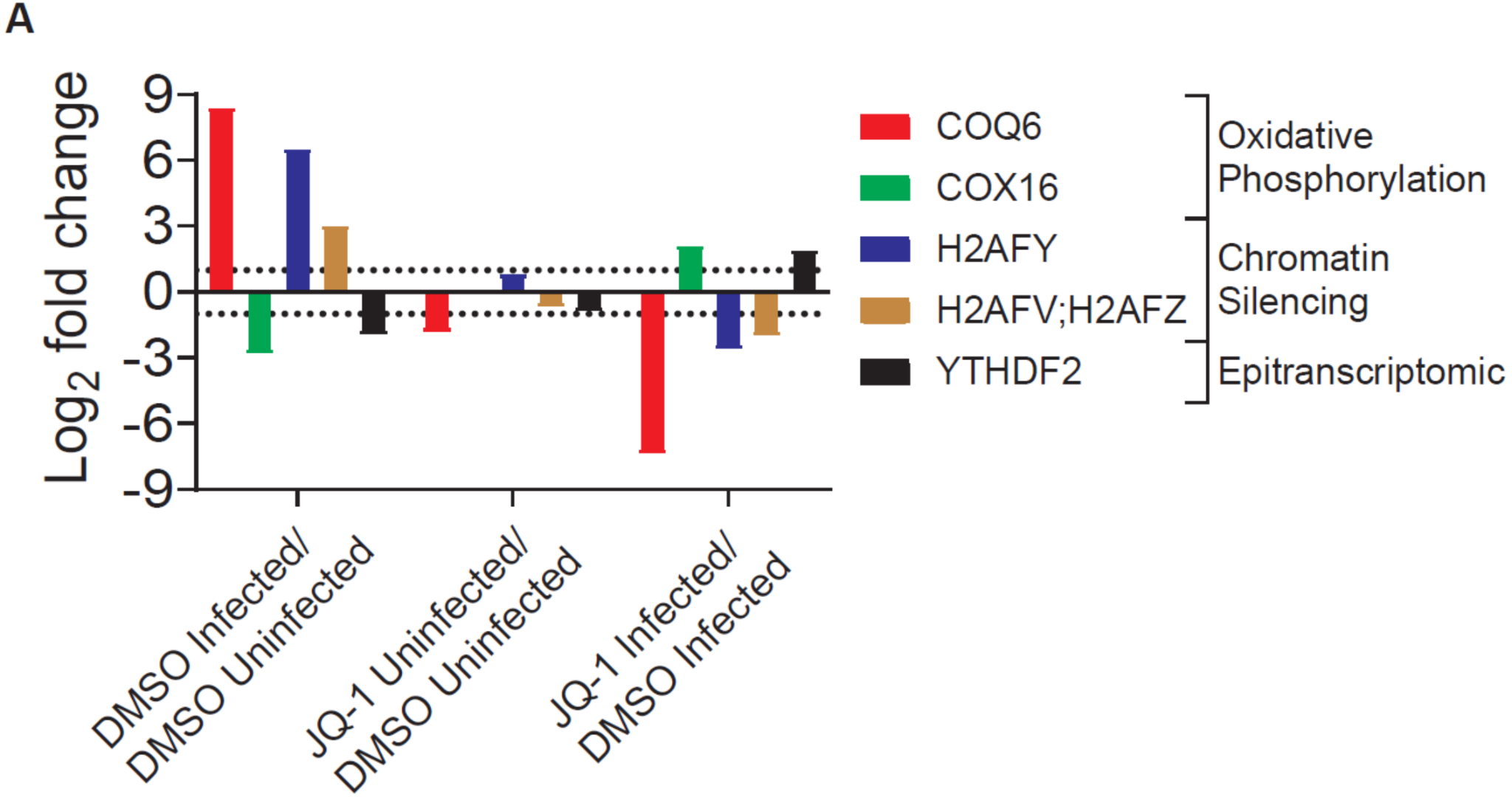

**Supplementary Figure 4. SARS-CoV-2 infection modulates expression of proteins involved in oxidative phosphorylation, chromatin condensation, and epitranscriptome in lung epithelial Calu-3 cells.**

(**A**) Log_2_FC analysis of differentially expressed proteins implicated in oxidative phosphorylation (COQ6 and COX16), chromatin silencing (H2AFY, H2AFV & H2AFZ), and epitranscriptomics (YTHDF2) from selected contrasts. The bars indicate relative log_2_FC in expression between experimental groups in each contrast, and proteins with a relative log_2_FC of 1 and a FDR≤0.05 were considered significant.

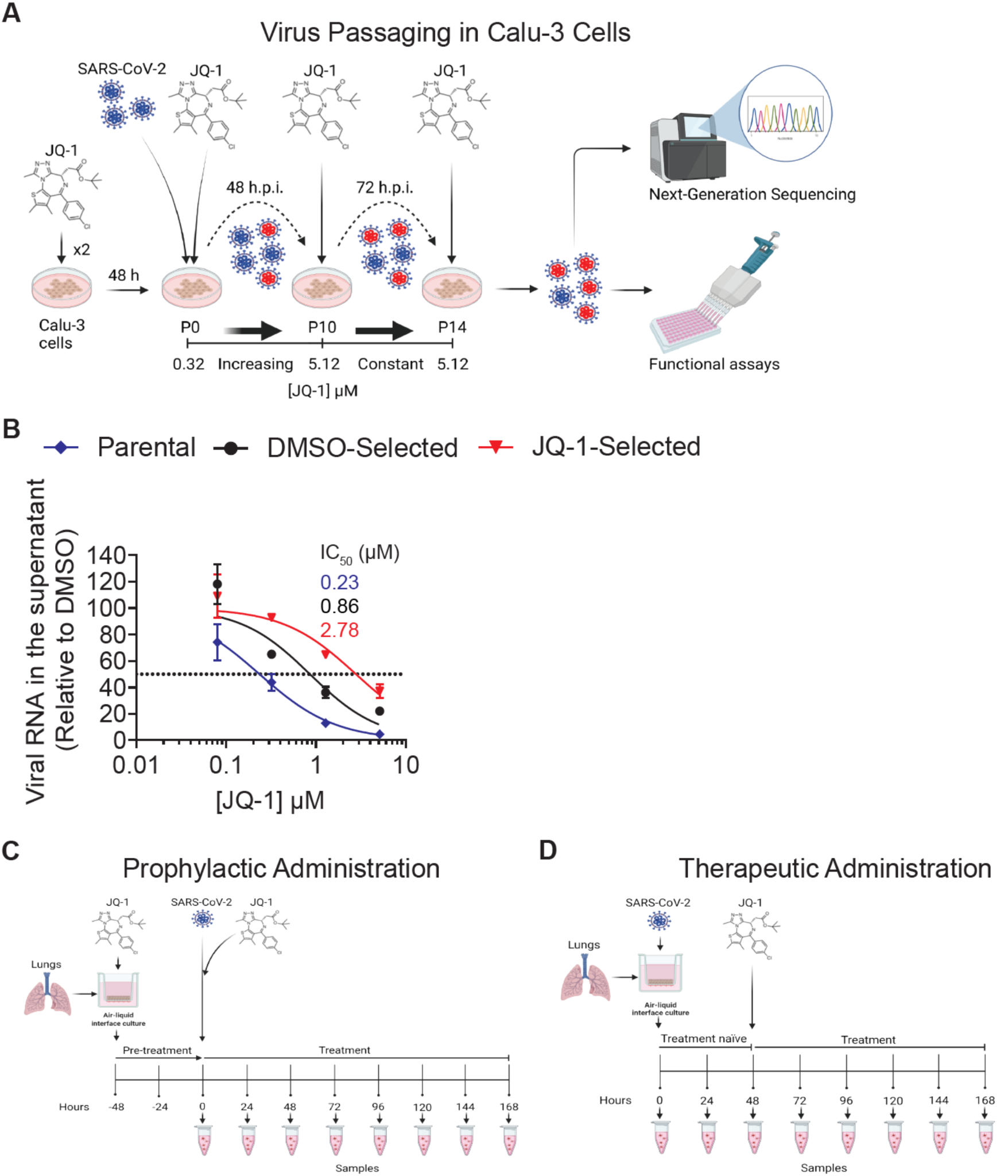

**Supplementary Figure 5. Serial passaging of SARS-CoV-2 under two-fold escalating concentrations of JQ-1 confers replication kinetic superiority under JQ-1 treatment in lung epithelial Calu-3 cells.**

(**A**) Schematic diagram of SARS-CoV-2 passaging in Calu-3 cells under two-fold escalating concentrations (0.32, 0.64, 1.28, 2.56, and 5.12 µM) of JQ-1. Each drug concentration was kept constant for two passages before escalation by two folds for the next two passages. Samples were taken at 48 h.p.i. until passage ten, at which point the highest non-toxic concentration of JQ-1 (5.12 µM) was reached. Serial passaging continued for the next five passages under a constant concentration of JQ-1 (5.12 µM), during which samples were taken at 72 h.p.i.

(**B**) JQ-1 dose-dependent inhibition curves determined in JQ-1-treated Calu-3 cells infected with parental and passaged (P14) SARS-CoV-2 virions (MOI 0.1) from four independent experiments.

Schematic diagrams of SARS-CoV-2 growth kinetics in hBAECs in the presence of DMSO or JQ-1 (2.56 µM). hBAECs were pre-treated twice for 48 hours with a drug, followed by SARS-CoV-2 (2x10^4^ PFUs) infection and drug administration to constitute a (**C**) prophylactic approach. hBAECs were followed for seven days, during which samples were taken after 24 hours followed by fresh drug administration. To constitute a (**D**) therapeutic approach, drug-naïve hBAECs were infected with SARS-CoV-2 (2x10^4^ PFUs) for 48 hours to establish production infection, followed by consecutive drug administration after 24 hours during which samples were taken until 168 hours post infection.

**Supplementary Table 1**. Significant enrichment of transcription factor binding motifs in the accessible ATAC-seq peaks in (**A**) DMSO-treated infected versus DMSO-treated uninfected, (**B**) JQ-1-treated uninfected versus DMSO-treated uninfected and (**C**) JQ-1-treated infected versus DMSO-treated infected contrasts. The motif search was conducted using the DREME algorithm to annotate the motifs to the known transcription factor families. The height of the letters represents the frequency of each base in the motif.

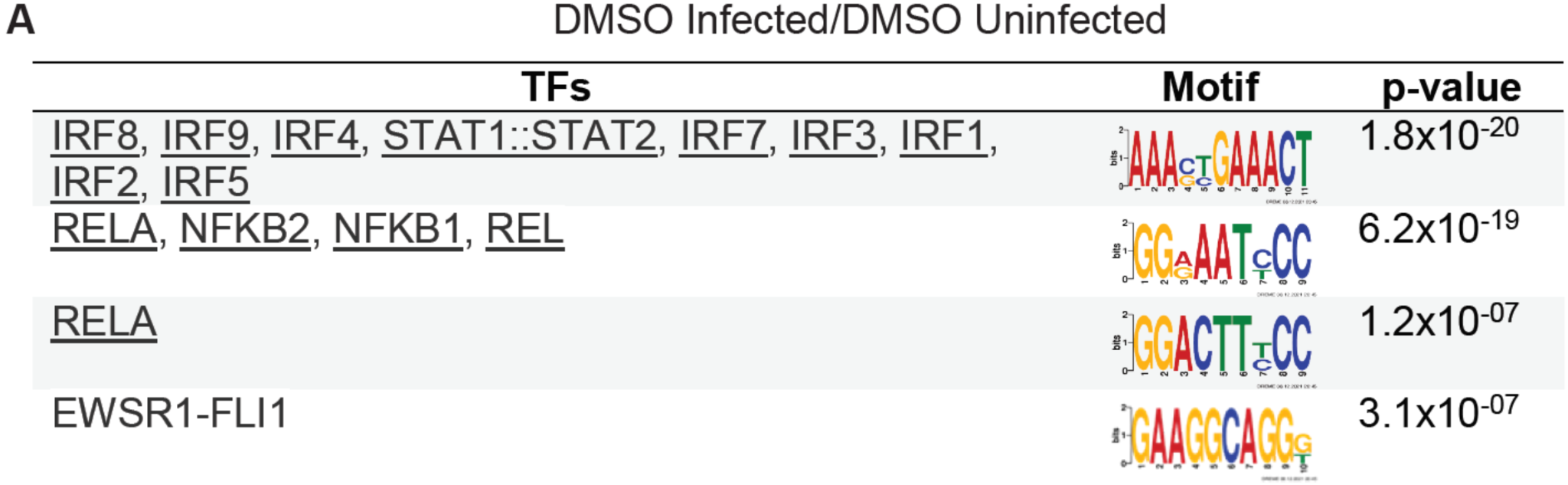

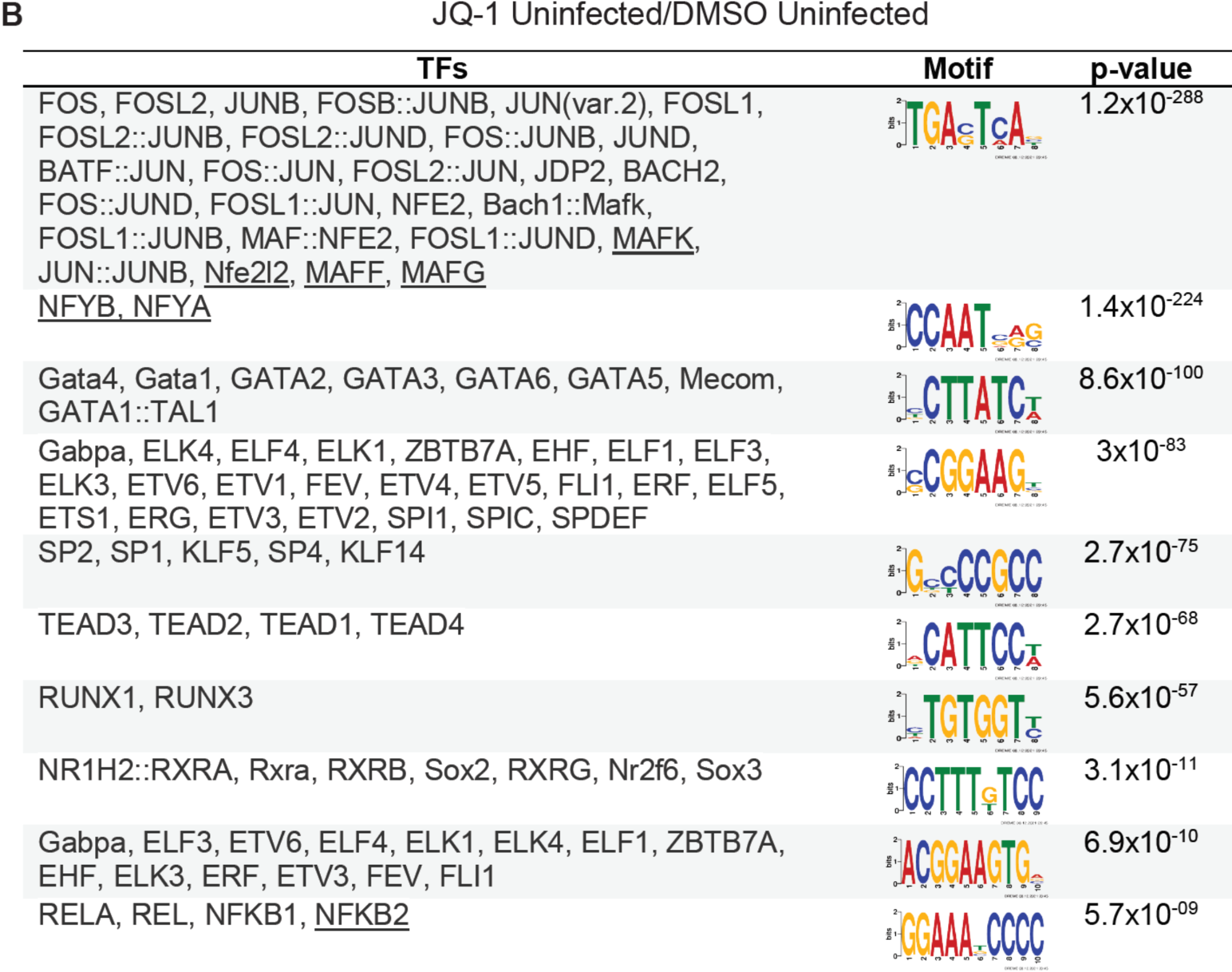

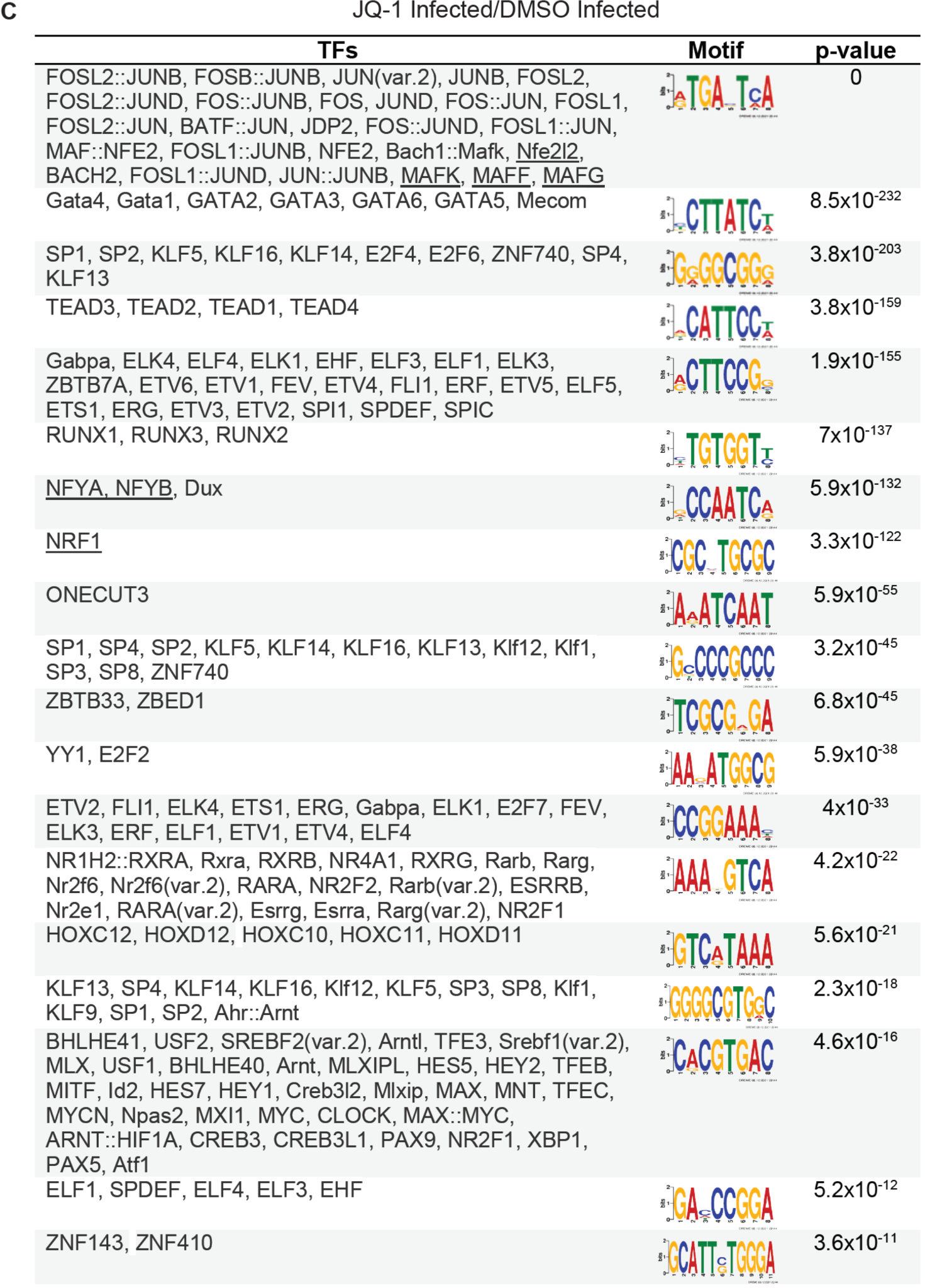
